# Spectrin condensates provide a nidus for assembling the periodic axonal structure

**DOI:** 10.1101/2024.06.05.597638

**Authors:** Nicholas P. Boyer, Rohan Sharma, Theresa Wiesner, Antoine Delamare, Florence Pelletier, Christophe Leterrier, Subhojit Roy

## Abstract

Coordinated assembly of individual components into higher-order structures is a defining theme in biology, but underlying principles are not well-understood. In neurons, α/β spectrins, adducin, and actinfilaments assemble into a lattice wrapping underneath the axonal plasma membrane, but mechanistic events leading to this periodic axonal structure (PAS) are unclear. Visualizing PAS components in axons as they develop, we found focal patches in distal axons containing spectrins and adducin (but sparse actin filaments) with biophysical properties reminiscent of biomolecular condensation. Overexpressing spectrin-repeats – constituents of α/β-spectrins – in heterologous cells triggered condensate formation, and preventing association of βII-spectrin with actin-filaments/membranes also facilitated condensation. Finally, overexpressing condensate-triggering spectrin repeats in neurons before PAS establishment disrupted the lattice, presumably by competing with innate assembly, supporting a functional role for biomolecular condensation. We propose a condensation-assembly model where PAS components form focal phase-separated condensates that eventually unfurl into a stable lattice-structure by associating with subplasmalemmal actin. By providing local ‘depots’ of assembly parts, biomolecular condensation may play a wider role in the construction of intricate cytoskeletal structures.

## INTRODUCTION

Super-resolution microscopy revealed a subplasmalemmal cytoskeletal lattice in axons, composed of α/β-spectrins, adducin, actin, and other associated proteins^1^. This periodic axonal structure (PAS) is thought to be a stable meshwork with parallel arrays of actin filaments spaced at ∼ 190 nm by spanning spectrin tetramers – cytoskeletal integrators composed of α/β-spectrins. Recent studies suggest that the actin may be organized as ‘braids’ twisting along the shaft of the axon^2–4^. The PAS has been seen in axons from a variety of species^5^, as well as in living neurons^6,7^ and brain slices^1,6^. Several functions have been attributed to the PAS, such as conferring structural stability to the axon^8–10^, acting as a scaffold for organizing signaling complexes^11,12^, and regulating axonal diameters^13,14^.

While substantial progress has been made in identifying component parts of the PAS, mechanistic events leading to the formation of these intricate structures are less clear. A unique aspect of PAS development is that local availability of the constituent proteins along the axon-shaft likely depends upon axonal transport from distant cell-bodies. Key constituents of the PAS such as spectrin, adducin and actin are known to move in slow axonal transport^15–20^, putting further constraints on the availability of building-blocks for assembly. How are all the necessary components for assembling the periodic lattice readily available in the axon shaft – particularly during periods of axonal growth – and how does the complex structure arise from these building blocks? Contrasting models have been proposed to explain the origin of the PAS. One study proposed a ‘propagation model’ where the PAS first emerges in the proximal axon adjacent to the cell body, followed by distal propagation of the lattice structure^21^. However, there is no direct evidence for this model, and no precedence for such templated proximal to distal assembly along the axon. A more recent study reported focal patches of spectrin in distal axon-shafts of developing neurons^22^ which is also inconsistent with the propagation model. The latter study proposes a different model where the PAS originates from the base of growth cones, with distal spectrin patches coalescing over time to form a continuous periodic structure. However, these conclusions were drawn from fixed images, and there is no evidence for such coalescence.

Revisiting the issue, we systematically visualized PAS components in cultured hippocampal neurons as they develop. We confirmed the observation that the PAS in the most proximal part of axons was established early, and focal patches of spectrins are present in distal growing axons. Examining these patches, we saw that they contained α/β-spectrins and adducin, but scant filamentous actin (F-actin). Collectively, our experiments in non-neuronal cells and neurons support the view that these distal spectrin/adducin patches are biomolecular condensates – driven by phase separation of spectrin-repeats – that associate with sub-plasmalemmal actin filaments to ultimately form the mature PAS. In our model, spectrin condensates act as transient depots for storing the building blocks necessary for PAS assembly, suggesting a role for phase separation in cellular homeostasis. We posit that such mechanisms may play a wider role in assembling cytoskeletal structures in other contexts.

## RESULTS

### Distal axons contain patches of spectrins and adducin that with sparse actin filaments

We first visualized the distribution of key PAS proteins – spectrins, adducin and actin – in developing axons of mouse cultured hippocampal neurons fixed between 2-10 days *in vitro* (DIV, **Fig. 1A**). Cultured hippocampal neurons develop in a stereotypical manner, with multiple tapering dendrites and a single elongated axon (**Fig. 1B**). The staining pattern of spectrins and adducin over time is shown in Figures 1 **C-E**. At early developmental stages (2 DIV), spectrins and adducin were concentrated in the proximal axon – defined here as the proximal third of its total length – near the presumptive axon initial segment (AIS, yellow bracket), with the staining tapering off distally (**Fig. 1C**). However, by 7 DIV, small, isolated patches of spectrin and adducin began to appear along distal axons, defined here as distal third of its total length (**Fig. 1D-E**). These patches were ∼1-4 µm in length – distinct from the proximal continuous segment of spectrin/adducin staining – and became more frequent and brighter as axons developed. To determine the quantitative distribution of these patches, we designed an algorithm to measure staining discontinuities (**Fig. 1F**, “discontinuity factor” δ – see methods for details). Briefly, δ is more positive in regions with distinct patches of fluorescence, and more negative in homogenous regions. Measuring δ along axons from the soma to the growth cone, we found that patches formed by αII/βII-spectrin and adducin were consistently more pronounced in distal axons, whereas βIII-tubulin (morphologic marker) was generally smoother throughout (**Fig. 1G**). Note that in general, the distal patchiness of spectrins/adducin in the axon is more pronounced at later developmental stages.

**Figure 1:**
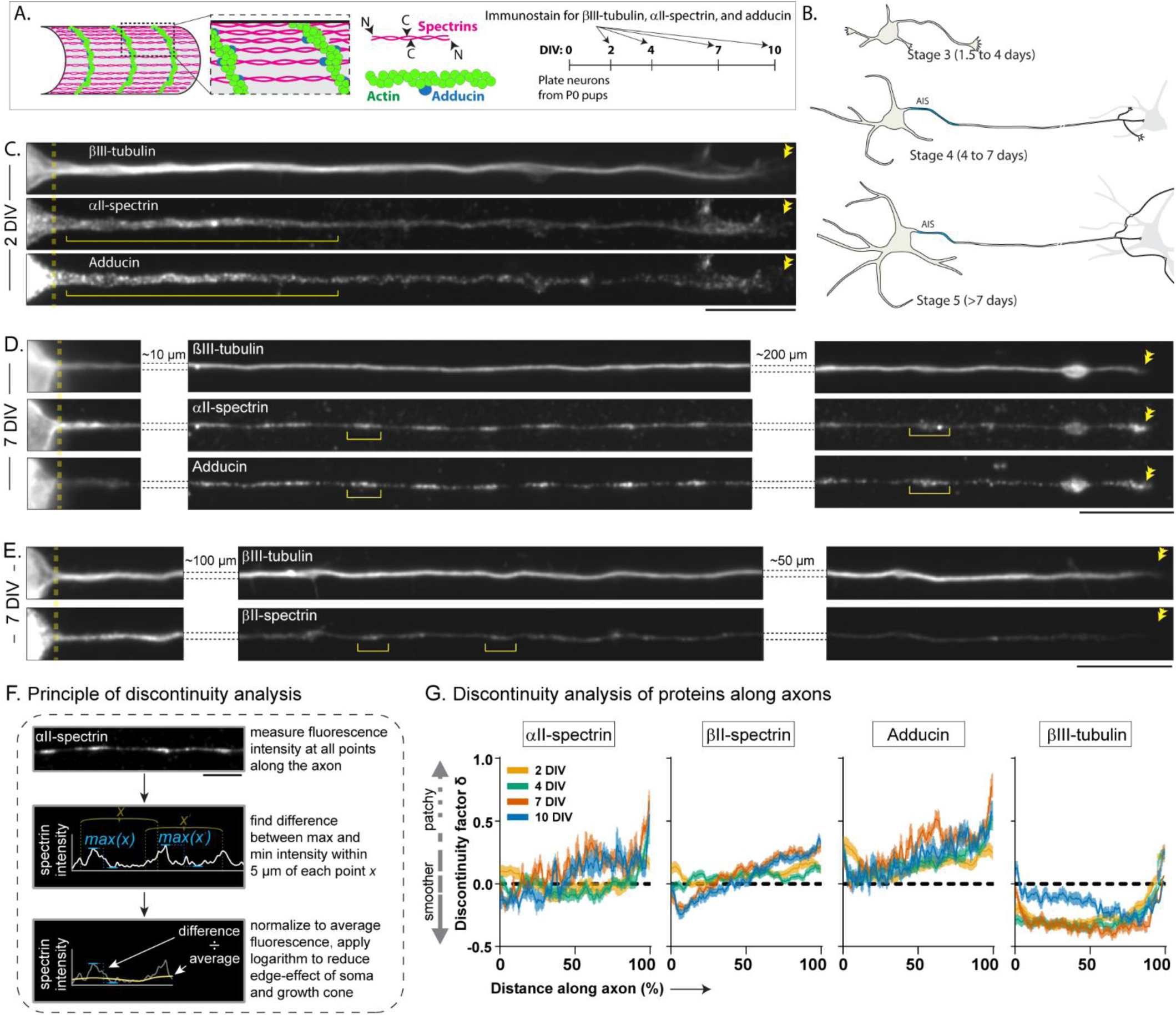
Spectrin/adducin patches in distal axon during development. **(A)** Schematic of the PAS and timing of experiments. Hippocampal neurons were obtained from P0 pups and cultured for 2, 4, 7 or 10 days in vitro (DIV) before fixation and immunostaining. **(B)** Schematic showing development of the axon through neuronal stages 3-5. Note formation of the axon initial segment (AIS) during stage 4 and synaptic contacts with target neurons at stage 5. **(C-E)** Representative images of 2 DIV and 7 DIV neurons stained for βIII-tubulin (volume marker), αII-spectrin or βII-spectrin, and adducin. Note accumulation of spectrin and adducin in the proximal axon. Soma/axon junction is marked by a dashed line, double arrowheads labels the growth cone, and a bracket marks patches along the axon. Scale bars = 10 µm. **(F)** Quantification of axon patches. Fluorescence intensity was measured at every point along the axon. The maximum and minimum fluorescence within 5 µm of each point was calculated, and the difference was normalized to the average fluorescence within 20 µm, giving an estimate of the discontinuity of fluorescence. The logarithm of this normalized difference was the discontinuity factor δ (see Methods for more details). Scale bar = 5 µm. **(G)** Discontinuity factors for αII/βII-spectrin, adducin and βIII-tubulin in neurons fixed at 2, 4, 7 or 10 DIV. Note that a higher positive δ indicates a patchier axonal distal axon, and in general, discontinuity in the distal axon increases as the axon develops (data from 9-46 cells for each time-point, from 3 independent cultures).

Next, we compared the distribution of axonal spectrin to actin. Towards this, we fixed and stained neurons for αII/βII-spectrin, filamentous actin (F-actin) and monomeric actin (G-actin), using βIII-tubulin staining as a morphological marker. The distribution of F-actin was discontinuous along the axon, and we noticed that distal spectrin patches often lacked high levels of F-actin (**Fig. 2A-B**). Quantitative analysis showed that significantly lessF-actin was associated with the spectrin patches found in the most distal parts of the axon (**Fig. 2C**, also see **Supp. Fig. 1A-C**). We also examined the nanostructure of these patches using stochastic optical reconstruction microscopy (STORM), at a timepoint when the spectrin/adducin patches are clearly visible in distal axons (6 DIV). As expected, proximal axons had a continuous periodic spectrin lattice with ∼190 nm periodicity; however, this structured organization was lost in distal axons, and imaging of patches revealed incomplete lattices and globular structures (**Fig. 2D**). Taken together, the data suggest that while the distal axonal patches are enriched in spectrins and adducin, they are often devoid of actin filaments, suggesting that the absence of F-actin may favor the assembly of these patches (also see βII-spectrin/actin data later).

**Figure 2:**
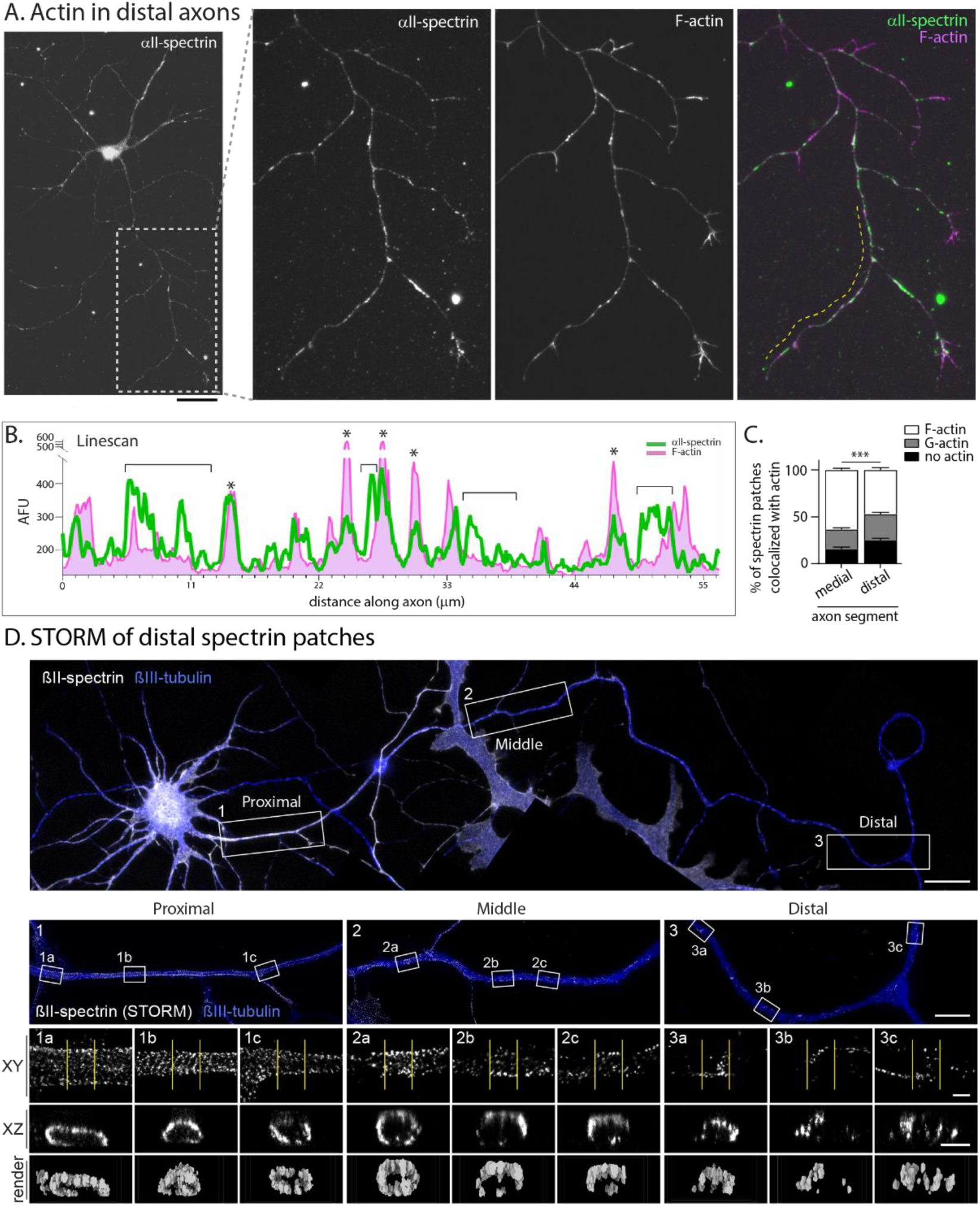
Paucity of actin filaments in distal spectrin patches. **(A)** Neurons at 7 DIV were fixed and stained for F-actin and αII-spectrin. In zoomed insets of distal axon, note that many spectrin patches lack F-actin, best seen in the merged image on right. Scale bar = 10 μm. **(B)** A maximum-intensity line-scan of the dashed yellow line from (A). Note that many patches of αII-spectrin do not coincide with peaks of F-actin (brackets), while some do (asterisks). **(C)** Quantification of αII-spectrin patch colocalization with F-actin (phalloidin) or G-actin (DNase). Note that more spectrin patches in the proximal and medial thirds of the axon contained F-actin, while patches in the distal third were more likely to colocalize with G-actin or contain no actin. Data from 31 cells from 3 independent cultures. **(D)** STORM imaging of βII-spectrin in proximal (1), middle (2), and distal (3) regions of a rat hippocampal neuron fixed at DIV 6. XY zooms and XZ axis projections of the regions delineated by yellow lines are shown below together with a 3D rendering of the section. Note that while proximal axons have the expected periodic appearance of spectrin, middle and distal axonal regions have interrupted periodicity with incomplete annular structures, or patchy distribution with no periodicity. Scale bars = 20 µm (top view), 5 µm (middle views), 500 nm (zooms and transverse sections).

### Spectrin patches in distal axons show condensate-like behavior

The morphology of spectrin/adducin patches in distal axons is reminiscent of membrane-less organelles or biomolecular condensates, which can range from small nanoclusters to “mesoscopic” protein condensates that are several hundred nanometers in size^23–26^. Specific to spectrin, previous studies in non-neuronal cells have shown that in response to DNA damage, nuclear αII-spectrin redistributes into bright foci resembling biomolecular condensates^27^. Components of biomolecular condensates can remain stably concentrated within a structure for long periods of time, but a cardinal feature is that molecules within such structures exchange with the surrounding cytoplasm within timescales of seconds to minutes^28,29^. To test if the axonal spectrin patches also behave in this manner, we performed fluorescence recovery after photobleaching (FRAP) experiments in cultured neurons (**Fig. 3A**). Neurons were transfected with low levels of αII-spectrin tagged to monomeric Green-Lantern (mGL), and the low expressors were selected for analyses (see Methods for details). While fluorescence was continuous in proximal regions, discrete patches of αII-spectrin:mGL were seen in distal axons (**Fig. 3B**), reminiscent of the pattern seen with endogenous αII-spectrin. These αII-spectrin:mGL patches were stable for the entire duration of imaging (15 minutes), and did not show any merging/splitting behaviors (**Supp. Movie 1**). FRAP experiments showed that while there was minimal recovery of αII-spectrin:mGL fluorescence in proximal axons – consistent with a stable lattice (see methods, **Supp. Fig. 2A**, and Zhong et al., 2019^21^) – fluorescence in photobleached distal αII-spectrin:mGL patches recovered partially within several minutes, indicating that cytosolic spectrin molecules from the axon shaft can exchange with the axonal patches. Studies have recognized phase separation in diverse cellular contexts, ranging from highly mobile granules that exchange rapidly with the surrounding cytosol^30,31^, to condensates that behave like solids, or have solid-like features^32,33^. Axonal αII-spectrin:mGL patches were also resistant to aliphatic alcohols that can disrupt the low-complexity interactions that drive some forms of phase separation (**Supp. Fig. 2B**); suggesting a more solid-like behavior, and also arguing against a model where the patches coalesce over time.

**Figure 3:**
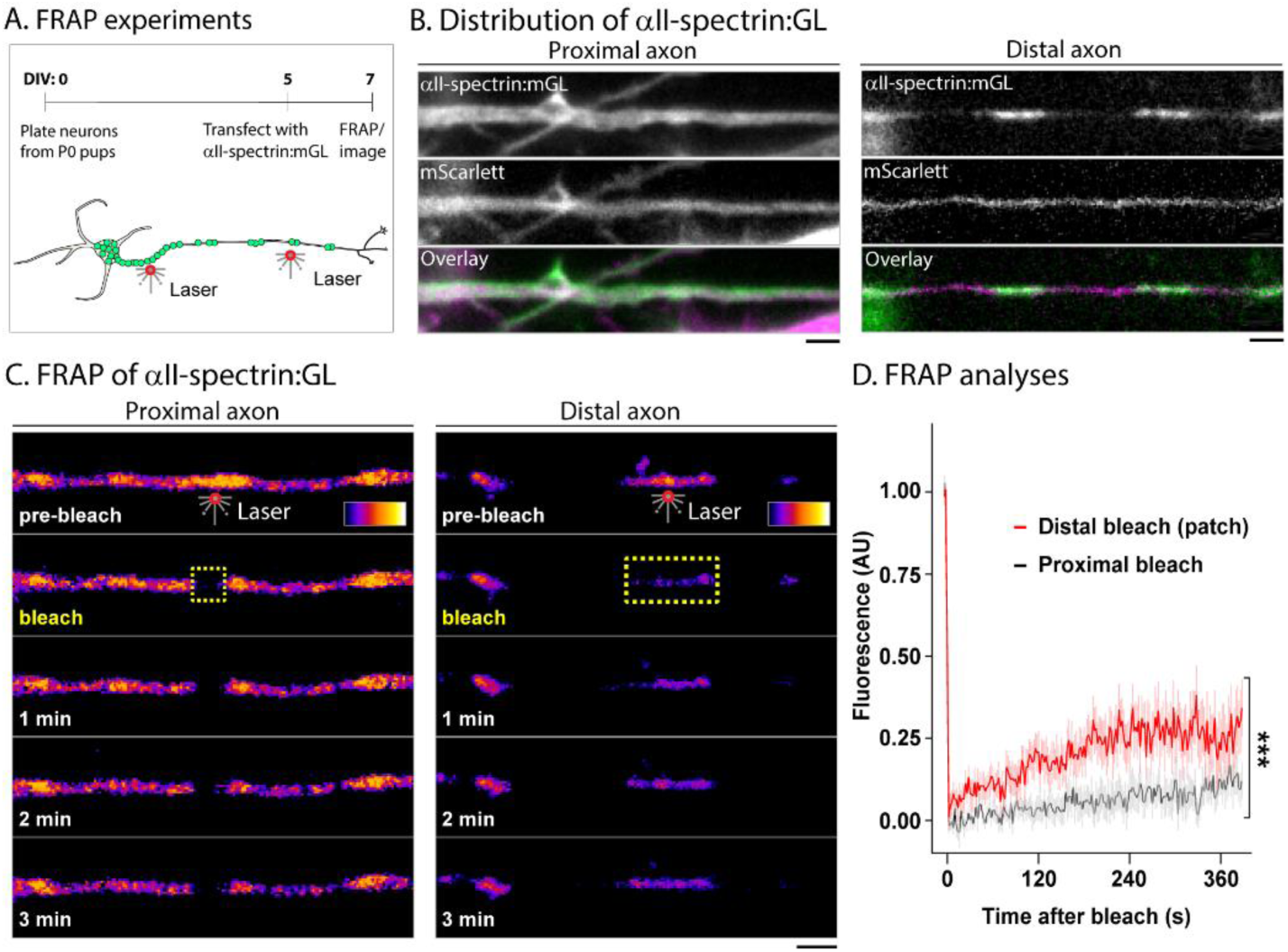
Spectrin in axonal patches can dynamically exchange with the axoplasm. **(A)** Schematic of live-cell imaging and FRAP experiments. 5 DIV neurons were co-transfected with αII-spectrin:mGL and mScarlet (volume filler). At 7 DIV, αII-spectrin:mGL was photobleached either in proximal axons (within 100 mm of soma), or in distal axonal patches. **(B)** Live imaging of αII-spectrin:mGL in proximal and distal axons. Note distinct patches of fluorescence in distal axon, reminiscent of the endogenous spectrin pattern. These patches were largely stationary over ∼15 minutes of imaging. Scale bars = 2 µm. **(C)** Timelapse images showing recovery of αII-spectrin:mGL fluorescence following photobleaching (yellow dashed region) in the proximal axon or distal patches. Note that while the proximal axon shows minimal recovery of fluorescence – consistent with a stable lattice – distal axonal patches partially recover within minutes. Scale bars = 2 µm. **(D)** Quantification of FRAP experiments showing significantly faster recovery in distal patches (n=12/condition, in axons from 3 independent cultures, two-way ANOVA, interaction ***p < 0.001).

### αII-spectrin can form bimolecular condensates in heterologous cells

A standard way of exploring phase separation is to overexpress putative condensate-components in heterologous cells, facilitating molecular interactions that would promote phase separation and generate focal-condensates^34–36^. Towards this, we transfected HEK293T cells with mGL-tagged αII/βII-spectrins or adducin and investigated putative condensate-like behaviors (**Fig. 4A**). The distributions of FL versions of mGL-tagged αII-spectrin, βII-spectrin, and adducin were distinct in HEK293T cells (**Fig. 4B**). While FL-αII-spectrin formed inclusions resembling condensates, FL-βII-spectrin – that has actin-binding domains^37^ – had a membranous pattern (also see **Supp. Movie 2**), and FL-adducin had a diffuse, cytosolic distribution (**Fig. 4B**). We noticed that cells with rounded inclusions sometimes also had large “islands” of fluorescence that were typically seen along the periphery of the cell, underneath the plasma membrane (**Supp Fig. 3**). Other studies on phase separation have also reported similar swaths of fluorescence upon overexpression of condensate-forming proteins^38,39^. While physical processes that control the size of condensates are not well understood^40,41^, it is known that native membraneless assemblies such as the nucleoli have large, non-normal variations in size-distributions^42,43^. Thus the behavior of spectrin condensates may be due to underlying biophysical processes that regulate the formation and maintenance of these assemblies, or alternatively, the large accumulations might be an effect of association of spectrins with the subplasmalemmal F-actin network, which alters the physical nature of these condensates. As controls, we also tested a synthetic membraneless organelle engineered from intrinsically disordered arginine/glycine-rich RGG domains^30^ that showed small focal inclusions in HEK293T cells, and soluble (untagged) mGL, which was diffusely distributed as expected (**Fig. 4C**).

**Figure 4:**
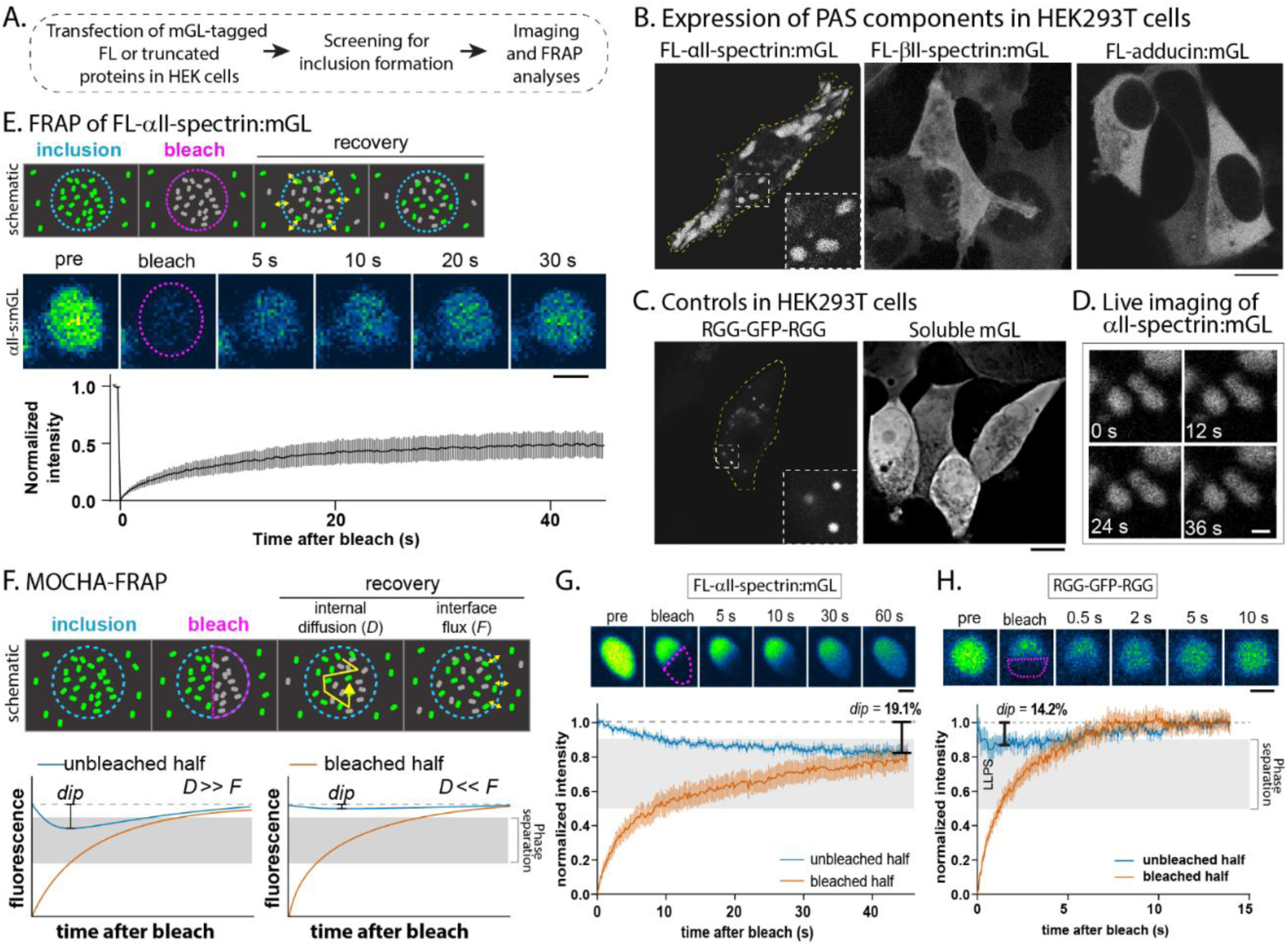
FL-αII-spectrin forms biomolecular condensates in heterologous cells. **(A)** Overall schematic of experiments expressing mGL-tagged FL or truncated αII/βII-spectrins/adducin in HEK293T cells. **(B)** Expression of mGL-tagged FL αII-spectrin, βII-spectrin and α-adducin in HEK293T cells. Note that FL-αII-spectrin formed distinct inclusions (zoomed in dashed inset), βII-spectrin showed a membranous localization (also see **Supp. Movie 2**), and α-adducin was diffuse. Scale bar = 10 µm. **(C)** Expression of a construct known to induce phase separation (engineered dimer of the RGG domain of yeast LAF1 protein fused to GFP), or soluble (untagged) mGL in HEK293T cells. Scale bar = 10 µm. **(D)** Frames from timelapse imaging of αII-spectrin:mGL inclusions in HEK293T cells show that they are largely stationary and do not undergo merging or splitting. Scale bar = 1 µm. **(E)** Top panel: Schematic showing kinetics of FRAP when a single inclusion is completely photobleached. Note that the recovery is due to diffusion of fluorescent molecules from unbleached regions into the bleached region, across the surface of liquid droplets (small yellow arrows). Middle/bottom panels: FRAP of αII-spectrin droplets shows recovery of 40-50% of fluorescence within 45 seconds (n=8, cells from 3 independent cultures, scale bar = 1 µm). **(F)** Top: Schematic showing the principal of MOCHA-FRAP experiments, in which one-half of a fluorescent inclusion is photobleached. Note that FRAP in this scenario is due to diffusion of fluorescent molecules from the unbleached region (long yellow arrow), as well as from across the surface of putative droplets (small yellow arrows). Below: Assuming that the fluorescence redistributes within the inclusion at diffusion rate *D* and fluorescence in the droplet as a whole recovers at interface flux-rate *F*, theoretical graphs show fluorescence of both unbleached and bleached droplet halves in a scenario in which *D* >> *F* (left), or when *D* << *F* (right). The depth of the fluorescence dip in the unbleached half correlates to the balance between these two rates, and is an indicator of liquid-like behavior. **(G)** MOCHA-FRAP analyses of αII-spectrin:mGL inclusions. Top panel: Representative FRAP images of a single αII-spectrin:mGL inclusion (top). The bleached region is marked with a dashed red line, note recovery of fluorescence in the bleached region. Bottom panel: Quantitative graphs of unbleached and bleached regions, obtained using the MOCHA-FRAP workflow. Black vertical bar indicates a dip depth of 19.1% at the location of minimum fluorescence of the unbleached half, which falls within the expected range for phase separation in the MOCHA-FRAP model (n=8, cells from 3 independent cultures). Scale bar = 1 µm. **(H)** MOCHA-FRAP experiments on RGG-GFP-RGG droplets which display a very fast internal redistribution and recovery of fluorescence, with a dip depth of 14.2% which is consistent with phase separation (n=9, cells from 3 independent cultures).Scale bar = 1 µm.

Live imaging of FL-αII-spectrin inclusions showed that they were stable and did not show significant merging/splitting behavior (**Fig. 4D** and **Supp. Movie 3**). Biomolecular condensation has two key features: 1) exchange of molecules between the condensate and surrounding cytosol; and 2) redistribution of molecules within the condensate by internal diffusion. Accordingly, we used two FRAP modalities to test if FL-αII-spectrin:mGL inclusions showed liquid-like behaviors. We first photobleached entire αII-spectrin:mGL inclusions in HEK293T cells (full-bleach) and evaluated FRAP (see schematic in **Fig. 4E**, top). As shown in the serial snapshots from a representative example and ensemble statistics (**Fig. 4E**, middle and bottom), fully-bleached αII-spectrin inclusions recovered approximately half of their fluorescence within ∼45 seconds, indicating that spectrin molecules can readily move across the inclusion surface. Next we used a recently described workflow called model-free calibrated half-FRAP (MOCHA-FRAP^32^) to evaluate redistribution of molecules within a single αII-spectrin:mGL puncta. The principle of this quantitative assay is that bleaching half of a condensate would ensure that recovery would only occur from the unbleached half – in addition to ongoing exchange between the cytoplasm and condensate – and the authors mathematically determined that a dip-depth of fluorescence between 10 and 50% would correspond to liquid-liquid phase separation (**Fig. 4F** – see Muzzopappa et al.^32^ for more details). MOCHA-FRAP analyses of FL-βII-spectrin puncta shows that the fluorescence dip-depth was 19.1% on average, which falls within the predicted range for phase separation (**Fig. 4G**). MOCHA-FRAP analyses of RGG-GFP-RGG condensates showed a fluorescence dip-depth of 14.2% – also within the predicted range for phase separation – while applying similar procedures on αII-spectrin:mGL inclusions in fixed cells did not show any change as expected (**Supp. Fig. 4A-B**).

### Spectrin repeats can form biomolecular condensates in heterologous cells

Next, we examined the expression of αII-spectrin fragments in HEK293T cells (**Fig. 5A**). FL-αII-spectrin is composed of 20 spectrin repeats, which are made of canonical coiled-coil domains that are found in all members of the spectrin superfamily of cytoskeletal integrators^44^. A portion of repeat 9 is homologous to SH3 domains, and EF-hands motifs at the C-terminus are involved in modulating interactions with βII-spectrins. As expected, algorithms predicting intrinsic disorder (see Methods) did not find significant disordered regions within FL-αII-spectrin (**Supp. Fig. 4C**), consistent with the highly structured coiled-coil domains that comprise spectrin repeats. Since multivalent interactions between SH3 domains are also known to participate in phase separation^24,33,40^, we generated a construct lacking this domain (ΔSH3). Additionally, recent studies show that coiled-coil domains can undergo phase separation. For instance, interactions between coiled-coil domains are known to drive the phase separation of several centrosomal proteins^45,46^, and many other proteins with coiled-coil domains are known to phase-separate, such as the structural Golgi protein GM130^47^, transcription factors and RNA-binding proteins^48,49^, endocytic proteins^50^, and others^51–53^. Even engineered proteins with coiled-coil domains can phase separate^54^, and simulations demonstrate that coiled-coil domains have a dramatically higher propensity for phase separation^55^. Accordingly, we also generated constructs containing variable number of spectrin repeats (encoding coiled-coil domains) but lacking EF-domains (15 or 9 repeats, see **Fig. 5A**)

**Figure 5:**
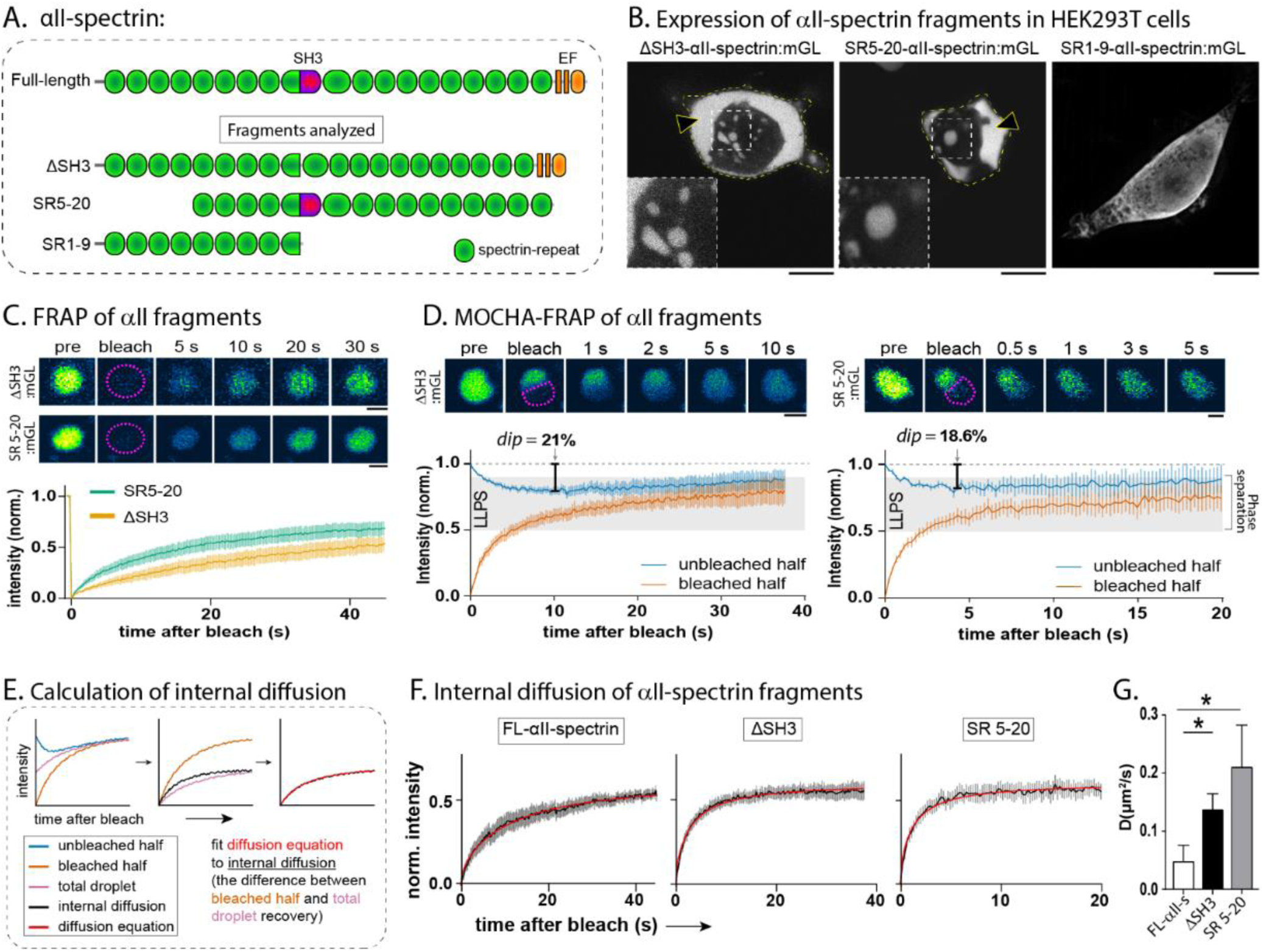
Spectrin-repeat domains within FL-αII-spectrin show droplet-like behavior. **(A)** Schematic showing FL and domain-deletion constructs of αII-spectrin that were tested for droplet formation in HEK293T cells. Constructs lacking the SH3 domain (ΔSH3), the N- and C-termini (SR5-20), or spectrin repeats 10-20 (SR1-9) were tagged with mGL. **(B)** Discrete, rounded inclusions were seen with ΔSH3 and SR5-20, but not with SR1-9 (highlighted in zoomed inset). Large swaths of fluorescence are also seen (black arrowheads), which may be related to extent of over-expression and biophysical properties (also see **Supp. Fig. 3** and Results, scale bars = 10 µm). **(C)** Full-inclusion FRAP analyses of mGL-tagged ΔSH3 and SR5-20 (top: images – dashed red line marks bleached region, bottom: FRAP-curves). Note that both αII-spectrin fragments show ∼ 40-60% fluorescence recovery (data obtained in cells from three independent cultures, scale bars = 1 µm). **(D)** MOCHA-FRAP analyses of mGL-tagged ΔSH3 and SR5-20 inclusions (top: images – dashed red line marks bleached region, bottom: fluorescence-curves from unbleached and bleached halves). Black bars indicate the dip-depths at the locations of minimum fluorescence of the unbleached half. Note that in both cases, the dip-depth falls within the predicted range for phase separation, highlighted in grey (n=7-10/condition, cells from 3 independent cultures, scale bars = 1 µm). **(E)** Schematic depicting steps for determining internal diffusion coefficient *D* from MOCHA-FRAP experiments. *D* is calculated from the Brownian diffusion curves (red) fit to the internal diffusion (black), which is equal to the fluorescence recovery of the bleached half (orange) with the total droplet recovery (pink) subtracted. **(F)** Graphs of internal diffusion from MOCHA-FRAP experiments with FL-αII-spectrin as well as SR5-20 and ΔSH3 constructs, with diffusion equation fits in red (n=7-10, cells from 3 independent cultures independent cultures). **(G)** Quantification of *D* from individual diffusion curves from populations of droplets summarized in (F). Deletion of the N/C-termini or SH3-domain results in faster internal diffusion than within full-length protein droplets. *, p < 0.05.

Eliminating the SH3 domain (ΔSH3) did not abolish the ability of αII-spectrin to form inclusions in this assay (**Fig. 5B**, left). Interestingly, overexpression of *isolated* spectrin-repeat domains (SR5-20) – without the EF-hands or repeats 1-4 that allow tetramerization with βII-spectrin – also formed inclusions in this experimental paradigm (**Fig. 5B**, middle), but a construct containing only the first 9 spectrin repeats (SR1-9) was diffusely distributed (**Fig. 5B**, right). Next, we tested putative condensate-like behaviors of these inclusions by FRAP and MOCHA-FRAP. Inclusions formed by both ΔSH3 and SR5-20 inclusions showed fluorescence recovery after full-bleaching, consistent with phase separation, though recovery of ΔSH3 was more restricted, perhaps due to the EF-domains (**Fig. 5C**). MOCHA-FRAP analyses of both ΔSH3 and SR5-20 showed that dip-depths for both deletion constructs were within the expected range for phase separation (**Fig. 5D**). MOCHA-FRAP also allows for the calculation of internal diffusion, as the difference between total droplet fluorescence recovery and recovery of the bleached half represents only movement of fluorescence within the droplet (**Fig. 5E**). We fit a standard Brownian diffusion equation (see Methods) to this difference and found that though deletion of the N- and C-termini or the SH3 domain did not fully ablate droplet formation, these deletions increased the internal diffusion rate of both ΔSH3 and SR5-20 constructs (**Fig. 5F-G**). Collectively, the data indicate that isolated spectrin-repeats (>9) can trigger condensate formation.

### Eliminating actin and membrane-binding domains of βII-spectrin facilitates condensate-formation

As FL-βII-spectrin did not readily exhibit phase separation when expressed in HEK293T cells, we next examined domain deletions of βII-spectrin. FL-βII-spectrin has calponin-homology actin-binding domains (CH-CH) at its N-terminus, followed by 17 spectrin repeats – consisting of coiled-coil domains – and a plasma-membrane binding pleckstrin homology (PH) domain at its C-terminus (**Fig. 6A**). Analyses of disorder-probability also suggested short putative disordered segments flanking the PH domain (**Supp. Fig. 4D**), prompting us to examine this segment as well. Unlike FL-βII-spectrin, that was enriched near the cell surface, striking inclusions were seen with overexpression of βII-spectrin fragments lacking the actin-binding (ΔCH-CH) domain, or a construct containing only spectrin-repeats (SR 1-17); while a fragment containing the PH-domain and predicted disordered regions was cytosolic (**Fig. 6B**). FRAP of SR1-17 inclusions showed substantial recovery after photobleaching – suggesting droplet-like behavior – but recovery for ΔCH-CH inclusions was much lower (**Fig. 6C**). MOCHA-FRAP analyses supported this observation, showing that only the dip-depth of SR 1-17 inclusions fell within the range of droplet-like behavior (**Fig. 6D**). Despite these differences in dip-depths, we did not find significant differences between internal diffusion rates of SR1-17 and ΔCH-CH inclusions (**Fig. 6E**). We also overexpressed fragments of adducin in HEK293T cells, but they either generated aggregates that did not behave like biomolecular condensates in FRAP assays, or were soluble like the FL protein (**Supp. Fig. 5**). Though the biological significance of these results is unclear, one possibility is that adducin molecules are passively recruited to spectrin condensates. In support of this, though FL-adducin was soluble in HEK293T cells when transfected alone (see **Fig. 3B**, right), it colocalized with αII-spectrin inclusions when co-transfected with FL-αII-spectrin (**Supp. Fig. 6**).

**Figure 6:**
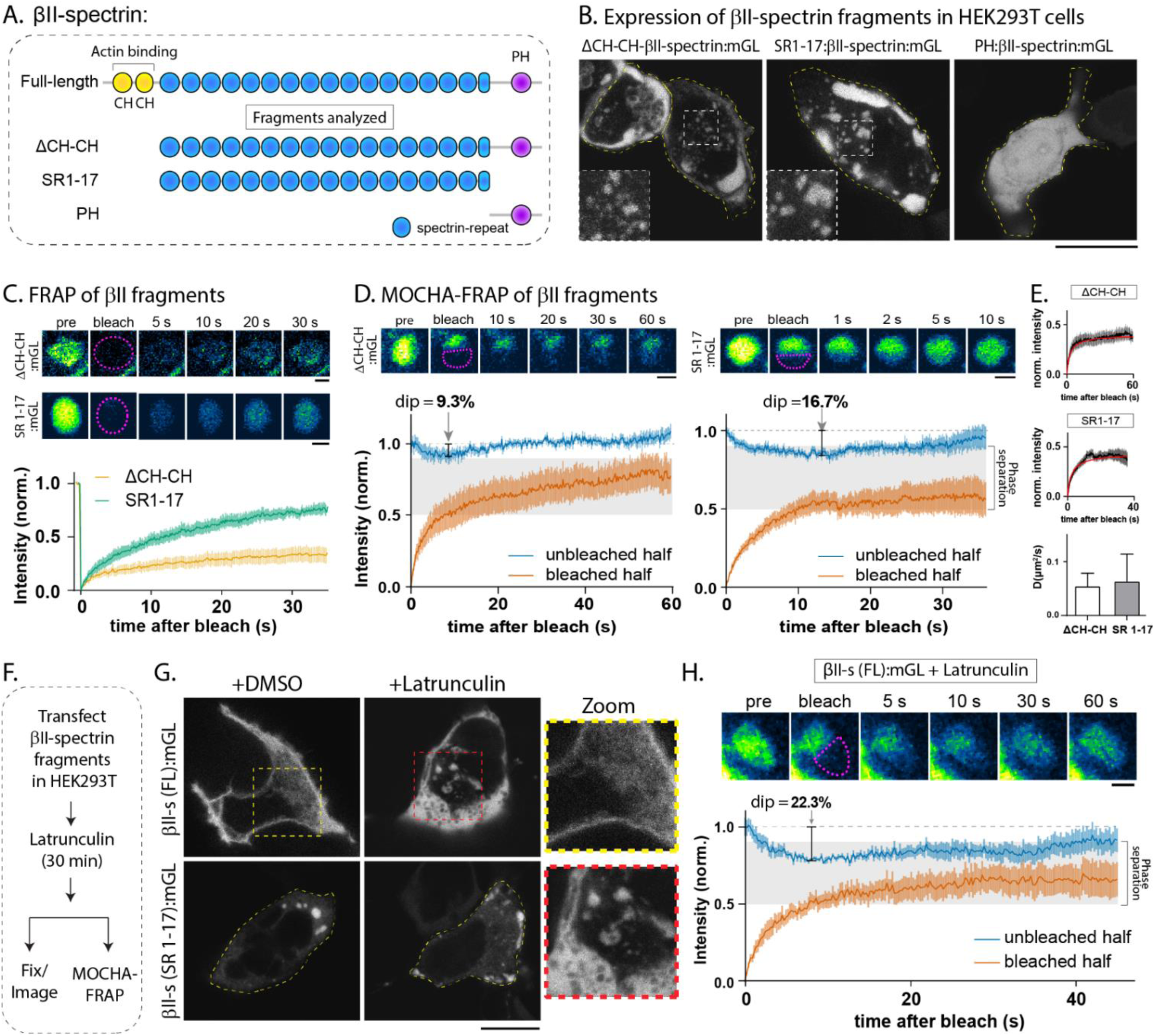
Isolated spectrin-repeats in FL-βII-spectrin can form condensates. **(A)** Schematic showing FL and domain-deletion constructs of βII-spectrin that were tested for droplet formation in HEK293T cells. Constructs lacking the actin-binding calponin homology domains (ΔCH-CH), containing only the 17 spectrin repeats (SR1-17), or containing only the C-terminal membrane-binding pleckstrin homology domain (PH) were tagged with mGL. **(B)** Discrete inclusions were seen with ΔCH-CH and SR1-17, but not with PH (highlighted in zoomed inset), suggesting that eliminating actin- and membrane-binding promoted inclusion-formation (scale bars = 10 µm). **(C)** Full-inclusion FRAP analyses of mGL-tagged ΔCH-CH and SR1-17 (top: images – dashed red line marks bleached region, bottom: FRAP-curves). Note that recovery was faster when both actin- and membrane-binding domains were deleted (SR1-17, compared to DCH-CH; n=7-9/condition, in cells from 3 independent cultures; scale bars = 1 µm). **(D)** MOCHA-FRAP analyses of mGL-tagged ΔCH-CH and SR1-17 inclusions (top: images – dashed red line marks bleached region, bottom: fluorescence-curves from unbleached and bleached halves). Black bars indicate the dip-depths at the locations of minimum fluorescence of the unbleached half. Note that only the dip-depth of SR1-17 falls within the predicted range for phase separation, highlighted in grey (n=7-9/condition, cells from 3 independent cultures, scale bars = 1 µm). **(E)** Internal diffusion graphs with fitted curves in red, and graphs of *D* calculated from population analyses (see method in Fig. 5E**-G**). Internal diffusion rates for ΔCH-CH and SR1-17 are similar, suggesting that MOCHA-FRAP differences may be driven by interface flux. Data from 7-9 droplets/cells from 3 independent cultures. Scale bars = 1 µm. **(F)** Testing the role of actin filaments in βII-spectrin droplet-formation. HEK293T cells transfected with FL-βII-spectrin or SR1-17-bII-spectrin were treated with 20 µM latrunculin A for 30 minutes to disrupt F-actin (or 4% DMSO as controls). **(G)** Top panels show that upon latrunculin treatment, there is redistribution of FL-βII-spectrin from its normal membranous distribution to intracellular inclusions (zoomed insets on right). Bottom panels show effects of latrunculin on SR1-17 inclusions. Representative images shown, scale bar = 10 µm. **(H)** MOCHA-FRAP analyses of FL-βII-spectrin inclusions formed after latrunculin-treatment. Top panel shows time-lapse images of an inclusion that was partially photobleached (bleached region highlighted by red dashed line), bottom panel shows curves from MOCHA-FRAP analyses. Note that the dip-depth (22.3%, black vertical bar) is within the expected range for phase separation (grey box; n=6 from 3 independent cultures; scale bar = 1 µm).

Taken together, quantitative analyses of βII-spectrin (FL and fragments) indicate that removing both the actin- and membrane-binding domains facilitates the formation of biomolecular condensates. Specifically, the isolated spectrin repeats of βII-spectrin showed the most robust evidence for phase separation. One prediction of this model is that preventing the association of FL-βII-spectrin with F-actin should also trigger condensate-formation. To test this, we transfected HEK293T cells with FL-βII-spectrin (or SR1-17 as a control), treated the cells with latrunculin to disrupt actin filaments, and examined the cells by imaging and MOCHA-FRAP (**Fig. 6F**). Interestingly, while FL-βII-spectrin was membranous in control cells as expected, treatment with latrunculin induced the formation of condensate-like structures in cells (**Fig. 6G**, top panels). Latrunculin treatment had little effect on pre-existing SR1-17 condensates, though there was a change in morphology, perhaps due to an increase in liquid-like behavior (**Fig. 6G**, bottom panels). MOCHA-FRAP of latrunculin-induced FL-βII-spectrin inclusions showed that the fluorescence dip-depth was within the range for phase separation (**Fig. 6H**).

### Early overexpression of condensate-forming spectrin repeats disrupts endogenous PAS assembly

Previous studies have shown that cytoskeleton-disrupting drugs can disassemble the PAS, but intrinsic structural determinants of PAS assembly are less clear. Our experiments advocate a model where α/β-spectrin repeats can organize into biomolecular condensates that form a nidus from which the PAS is ultimately assembled. If so, we reasoned that overexpression of condensate-forming spectrin-repeat fragments in neurons – before the PAS is fully assembled – would interfere with putative endogenous phase separation mediated by spectrins, leading to a disruption of the PAS. However, following this logic, overexpression of the same fragments after the establishment of the PAS would have little effect in the short-term, due to the known stability of the mature lattice structure^19^. To test this idea, we overexpressed condensate-forming βII-spectrin-repeat fragment (SR1-17, tagged to mScarlet) in cultured hippocampal neurons during early PAS assembly (from 3 to 6 DIV) or once it is fully formed along the proximal axon (from 7 to 10 DIV), and examined periodicity of the PAS within 100 µm from the soma using structured illumination microscopy (**Fig. 7A**). Note that the antibody in these experiments to visualize endogenous βII-spectrin recognizes an epitope at the C-terminus of the protein and is not expected to see the over-expressed SR1-17 fragment. Expression of SR1-17:mScarlet between 3 and 6 DIV disrupted the PAS seen in proximal axons, whereas expression of soluble mScarlet had no effect (**Fig. 7B-C**). On the other hand, when SR1-17:mScarlet was overexpressed between 7 and 10 DIV – a time when the PAS is expected to be established in the proximal axon – it had no effect on the periodic structure (**Fig. 7D-E**).

**Figure 7:**
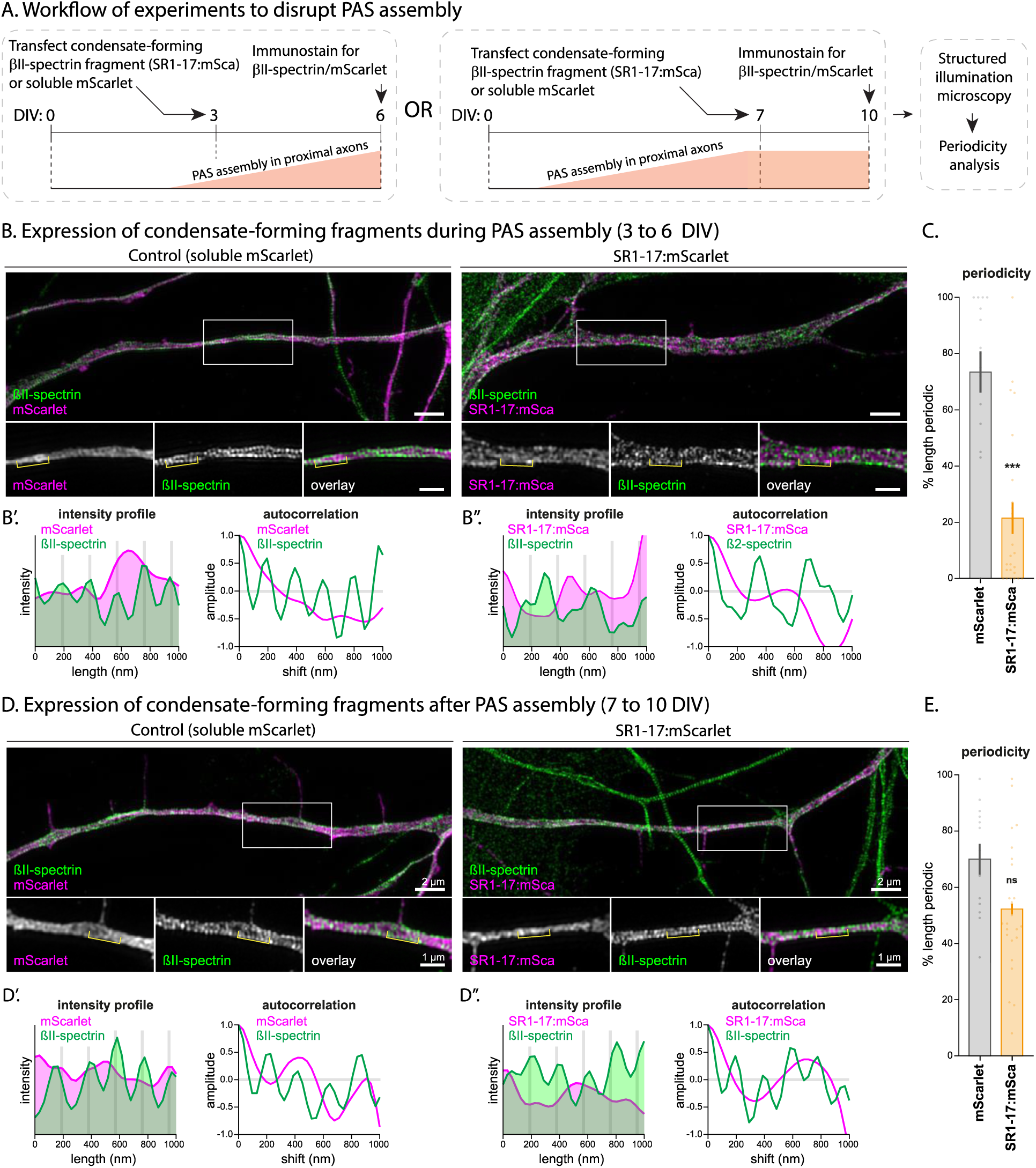
Early overexpression of condensate-forming spectrin repeats disrupts endogenous PAS assembly in neurons. **(A)** Overall experimental design to test if condensate-forming spectrin repeats can act in a dominant-negative manner to disrupt PAS assembly. Spectrin repeats of βII-spectrin (SR1-17:mScarlet) or soluble mScarlet were transfected in cultured hippocampal neurons either during (3 to 6 DIV), or after (7 to 10 DIV) establishment of the endogenous PAS in the proximal axon (region within 100 mm from soma). Neurons were subsequently fixed, stained with an antibody against the C-terminus of βII-spectrin (that would label endogenous βII-spectrin but not the transfected SR1-17), and spectrin-periodicity in axons was evaluated by SIM. **(B)** Periodicity in proximal axons of neurons transfected with SR1-17:mScarlet (or soluble mScarlet) at 3 DIV. Note that overexpression of SR1-17 impeded the assembly of the PAS; zoomed insets below (scale bars = 2 µm for large images, 1 µm for zoomed insets). **(B’,B”)** Fluorescence intensity profiles and autocorrelation analysis of axonal segments from **(B)** – zoomed insets, analyzed region marked by white brackets. Note disrupted periodicity upon transfection with SR1-17. **(C)** Quantification of the βII-spectrin periodicity in proximal axons at 6 DIV (transfected at 3 DIV). Periodicity in neurons expressing SR1-17:mScarlet was significantly lower than in neurons expressing soluble mScarlet (data from 15 to 25 neurons from 3 independent cultures. ***, p < 0.001). **(D)** Periodicity in proximal axons of neurons transfected with SR1-17:mScarlet (or soluble mScarlet) at 7 DIV – zoomed insets below (scale bars = 2 µm for large images, 1 µm for zoomed insets). Note that overexpression of SR1-17after the assembly of the PAS has no effect on periodicity, suggesting that the over-expressed fragments interfere with the assembly, but not maintenance of the PAS. **(D’,D”)** Fluorescence intensity profile and autocorrelation analysis of axonal segments from **(D)** – zoomed insets, analyzed region marked by white brackets. Note no disruption of periodicity upon transfection with SR1-17. **(E)** Quantification of the βII-spectrin periodicity in proximal axons at 10 DIV (transfected at 7 DIV). Note that overexpression of SR1-17 at a later stage had no effect on periodicity (data from 13 to 24 neurons from 3 independent cultures., ns = not significant).

## DISCUSSION

We started these experiments to find a plausible explanation on how the PAS attains its intricate fully assembled periodic lattice in axons. In line with previous studies^6,21^, we found that as neurons grow, the PAS first assembles in the most proximal part of the axon, and the timing of this assembly coincides with the formation of the AIS. However, as axons develop, we also saw focal patches in distal axons that contained spectrins and adducin, but had low levels of F-actin. Super-resolution microscopy showed that these distal patches do not have an organized lattice, and although they appear stable by live imaging, FRAP experiments showed that molecules within these patches can exchange with the surrounding axonal cytoplasm – a behavior consistent with phase separation. Expression of FL PAS-constituent proteins and domain-fragments in heterologous cells collectively indicate that spectrin fragments from both α/β -spectrins can form biomolecular condensates. These experiments also showed that preventing association of βII-spectrin with actin filaments and membranes triggered the formation of condensates, suggesting that the presence or absence of F-actin in the axon could regulate condensate-like behavior of the PAS components and help assemble the periodic lattice. Finally, overexpressing condensate-triggering spectrin repeats early in neuronal development – during establishment of the PAS – disrupted its periodicity in the proximal axon, suggesting dominant-negative suppression of assembly by condensate-forming spectrin repeats incapable of binding actin.

### A Condensation-Assembly model for building the periodic axonal cytoskeleton

Previous studies have proposed a propagation model where the lattice initially assembles next to the cell-body/axon junction, and then propagates distally along the length of the axon^21^. Another study observed βII-spectrin patches in distal growing axons but did not evaluate these structures further; proposing a model where the PAS originates at the axon-tip, with distal patches coalescing over time to form a continuous organized lattice^22^. However, there is no direct evidence to support these contrasting models. Continuous, templated assembly of the PAS towards the distal axon – as predicted by the propagation model – or coalescence of the distal spectrin patches into an assembled periodic lattice has not been observed. Our experiments advocate a new model where biomolecular condensation and actin filaments play roles in the biogenesis and assembly of the PAS respectively.

In line with previous observations^6,21,22^, we found that as axons begin to grow, the PAS is first assembled in the most proximal regions. Though the mechanistic events underlying this initial assembly are unclear, it may relate to proximity of the initial segment to the soma, where components of the PAS are synthesized. At a time when the PAS is assembled near the soma-axon junction, distal axons have discrete patches containing spectrins and adducin, and we propose that the patches act as local depots for supplying the building blocks necessary to assemble the PAS. What are the mechanisms underlying the assembly of these axonal patches? Taken together, our experiments in neurons and non-neuronal cells strongly suggest that the spectrin repeats within α/β-spectrins form biomolecular condensates that provide the necessary environment for concentrating components of the PAS into distal axonal patches. While many biomolecular condensates are dynamic and show dripping and fusing behavior^40,54,56,57^, axonal patches are largely static and do not coalesce over a timescale of minutes (**Supp. Movie 1**), but exchange molecules with the surrounding cytoplasm (**Fig. 3C-D**) – consistent with known behaviors of solid-like condensates^33,58,59^. The solid-like behavior of spectrin inclusions – both in neurons and HEK293T cells (see **Figs. 4-6**) – was unexpected, but may reflect a need to place PAS building-blocks at precise positions along the growing axon. Notably, solid phases are recognized in many native organelles that are thought to assemble by phase separation^40,60,61^.

How are components from the axonal patches eventually integrated into a mature periodic lattice? Though we do not have complete answers to this difficult question, two observations suggest that actin filaments may play a role. First, spectrin/adducin patches tend to form in axonal regions that have relatively less F-actin (**Fig. 2**), suggesting that the paucity of actin filaments may be driving force for condensation in this setting. Second, while βII-spectrin:mGL largely localizes to the plasma membrane in HEK293T cells, deleting actin- and membrane-binding domains of βII-spectrin triggered phase separation (**Fig. 6**). Taken together, these observations are consistent with a scenario where gradual accumulation of actin filaments in the distal axon would translocate spectrins and adducin – components of the PAS that bind to each other – from the axonal patches to the subplasmalemmal network of actin filaments, where they can be assembled into the periodic lattice. Future studies using biophysical modeling, artificial reconstitution of the lattice, or *in vitro* phase separation assays with recombinant proteins may give a more detailed view of these events. It is also interesting to consider that the lack of a rigid lattice in the distal axons may be an evolutionary adaptation to allow plasticity of the distal axon and growth cone during development.

### Spectrin repeats as novel mediators of biomolecular condensation

Recent studies suggest that formation of the centrosome and central spindle relies on biomolecular condensation of coiled-coil proteins^46^, and many of the component proteins such as pericentrin, spd-5, centrosomin, and MAP65/PRC1 can undergo phase separation^33,62–64^. Several other proteins known to undergo phase separation have coiled-coil domains^47–53^, and proteins engineered to contain coiled-coil domains can be coaxed to form condensates^41^. Simulations show that as few as two coiled-coil domains can phase separate, and that the propensity to condensate scales up with the number of coiled-coil domains^55^. We found that isolated spectrin-repeat domains of α/β-spectrins can undergo phase separation in HEK293T cells (**Figs. 5-6**). Spectrin repeats fold as coiled-coil helices, and this repeating structure is a defining feature of the spectrin superfamily that helps different isoforms of spectrin bind to each other^44^. Our data show that at high protein concentrations – such as conditions in the HEK293T cells after overexpression – coiled-coil domains of the same spectrin isoform can presumably self-associate into biomolecular condensates. Though it is unclear how such high concentrations of spectrins are achieved inside the axon, one possibility is that this is a combined effect of: a) the potential for FL-αII-spectrin to form condensates, and b) a paucity of actin filaments in distal axons that increases the propensity of FL-βII-spectrin to undergo phase separation (due to spectrin-repeats within its sequence). Since α/β-spectrins are typically present together as heterotetramers, presumably this would create foci with high concentration of spectrin molecules. Regardless, our experiments add spectrins to the growing list of coiled-coil domain proteins that can undergo phase separation, and advocate a role for this process in the early assembly of higher-order cytoskeletal structures.

## Acknowledgements

This work was supported by an NIH grant to S.R. (R01NS075233), and a NINDS P30NS047101 grant to the UCSD microscopy core. C.L. acknowledges funding from the Agence National de la Recherche (ANR-20-CE13-0024), and INP NCIS imaging facility and Nikon Center of Excellence for Neuro-NanoImaging for service and expertise, with funding from Excellence Initiative of Aix-Marseille University,A*MIDEX, a French ‘Investissements d’Avenir’ program (AMX-19-IET-002) through the Marseille Imaging and NeuroMarseille.

## METHODS

### Animals

Primary mouse hippocampal neuron cultures were prepared from CD1 pups obtained from Charles River Laboratories (Cat#022-CD1). Rat neuron cultures were produced by extracting hippocampi from E18 rat pups from pregnant female Wistar rats (Janvier labs). All procedures were in agreement with the guidelines established by the European Animal Care and Use Committee (86/609/CEE) or guidelines established by the University of California San Diego IACUC Office.

### Plasmids, antibodies, and reagents

The pcDNA3.1-mGreenLantern (#161912) and pcDNA_RGG-GFP-RGG (#124939) plasmids were acquired from Addgene. All mGreenLantern-tagged constructs consist of ORFs inserted into the multiple cloning site of pcDNA3.1-mGreenLantern using a Gibson cloning kit (NEBuilder HiFi DNA Assembly Cloning Kit, New England Biolabs). Intersectin(SH3_5_) was cloned from a construct generously gifted by the lab of Dr. Pietro De Camilli (Yale School of Medicine). Murine βII-spectrin and human αII-spectrin constructs were cloned from templates generously gifted by the lab of Dr. Matthew Rasband (Baylor College of Medicine). βII-spectrin full-length, ΔCH_2_ (aa303-2363) and SR1-17 (aa303-2084), and αII-spectrin SR8-10 (aa783-1242) and SR1-9 (aa1-969) inserts were generated by PCR from templates. αII-spectrin full length protein, ΔSH3 (aa967-1026 replaced with GGGG), and SR5-20 (aa468-2310) constructs were generated from these templates by a commercial cloning service (VectorBuilder Inc.). Inserts for βII-spectrin IDR-PHD-IDR (aa2085-2363), full-length murine Adducin1 (Genbank NM_001024458), Adducin1 aldolase (aa141-332) and NES-IDR (aa377-391 fused to aa544-735) were generated by custom oligonucleotide synthesis (gBlocks Gene Synthesis, Integrated DNA Technologies, Inc.). Adducin1 ΔIDR (aa1-577) and ΔN-term (aa141-735) inserts were cloned from Adducin1:mGreenLantern by PCR. Scarlet-tagged constructs were created by Gibson assembly of inserts generated by PCR from βII-spectrin SR1-17:mGreenLantern and αII-spectrin SR8-10:mGreenLantern inserted into pCCL-mScarlet (Addgene, #209889).

Primary antibodies used in this study: rabbit polyclonal against Adducin1 (Abcam ab51130), mouse monoclonal against αII-spectrin (Abcam ab11755), mouse monoclonal against βII-spectrin (BD Bioscience #612563), chicken polyclonal against βIII-tubulin (Abcam ab41489), and chicken monoclonal against mScarlet (Synaptic systems #409006). Secondary antibodies used in this study: goat anti-chicken Alexa Fluor 647 conjugate (Invitrogen A21449), goat anti-rabbit Alexa Fluor 594 conjugate (Invitrogen A11012), and goat anti-mouse Alexa Fluor 488 conjugate (Invitrogen A11001).

Additional reagents used in these studies: Alexa Fluor 488 conjugated phalloidin (Invitrogen A12379), Alexa Fluor 594 conjugated Deoxyribonuclease I (Invitrogen D12372), Latrunculin A (Calbiochem #428026), and DMSO (Sigma-Aldrich D2650).

### Primary Cell culture

Mouse hippocampal neurons were obtained from brains of postnatal (P0) CD-1 mice and plated on Cellvis glass-bottom dishes as described previously^65,66^, in compliance with University of California guidelines. Briefly, Cellvis dishes were coated with 100μL of 1 mg/mL poly-D-lysine + 1 mg/mL mouse laminin overnight at room temperature, washed thrice with ddH_2_O, and air-dried before plating. Hippocampi from P0 pups were dissected in ice-cold dissection buffer (HBSS, 4.44 mM D-glucose, and 6.98 mM HEPES) and incubated in 0.25% Trypsin-EDTA at 37°C for 15 minutes. Neurons were then dissociated in plating media (90% Neurobasal/B27 + 10% FBS, Life Technologies) by trituration. Neurons were plated at a density of 3,000 cells/100μL (immunostaining assays) or 30,000 cells/100μL (FRAP assays) of plating media. Neurons were maintained in NB27 media (Neurobasal media supplemented with 2% B27 and 1% GlutaMAX) in an incubator at 37°C and 5% CO_2_.

Rat hippocampal neurons were obtained from E18 rat pups from pregnant female Wistar rats. Hippocampi were dissected and homogenized by trypsin treatment followed by mechanical trituration and seeded on 18-mm diameter round, #1.5H coverslips at a density of 12,000-20,000 cells/cm^2^ for 3 hours in serum-containing plating medium (MEM with 10% fetal bovine serum, 0.6% added glucose, 0.08 mg/mL sodium pyruvate, 100 UI/mL penicillin-streptomycin). Coverslips were then transferred, cells down, to petri dishes containing confluent glia cultures conditioned in NB27+ medium (Neurobasal medium supplemented with 2% B-27, 100UI/mL penicillin/streptomycin and 2.5 μg/mL amphotericin) and cultured in these dishes (Banker method^67^).

### Immunostaining

Neurons were fixed in 4% paraformaldehyde in PEMS (80 mM PIPES, 5 mM EGTA, 2 mM MgCl_2_, 4% sucrose, pH 6.8) for 15 min at room temperature, followed by 3 rinses in 1X PBS. Cells were permeabilized in 0.2% Triton X-100 for 10 min, and blocked in 1X PBS containing 10% bovine serum albumin for 2 hours at room temperature. Neurons were incubated overnight at 4°C with primary antibodies in blocking buffer, washed three times with 1X PBS, then incubated with secondary antibodies labeled with Alexa Fluor fluorophores for 1 hour at room temperature. Cells were washed three times with 1X PBS and then mounted with Fluoro-Gel mounting media (Electron Microscopy Sciences #50-247-04). Immunostained neurons were imaged on a Nikon Eclipse Ti-E inverted epifluorescence microscope equipped with CFI Plan Apochromat VC 100X oil (NA 1.4; Nikon) objective, and an electron-multiplying charge-coupled device camera (QuantEM:512SC; Photometrics).

### Discontinuity Factor

The discontinuity factor was calculated from fluorescence linescans of full axons. This factor at a given point *x* along the axon was defined as the difference between the maximum and minimum fluorescence within 5 microns of *x*, normalized to the median fluorescence within 20 microns. This is expressed as the equation:

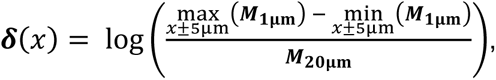

where ***M*_1μm_** and ***M*_20μm_** are the 1μm and 20μm micron moving medians, respectively. The 1 micron moving median was used to reduce the impact of visual noise on measurements. A logarithm was applied to the value in order to reduce edge effects from the increased intensity of the soma and growth cone.

### STORM Super-resolution microscopy

Rat hippocampal neurons from E18 embryos were cultured on 18 mm coverslips at a density of 6,000/cm^2^ following guidelines established by the European Animal Care and Use Committee (86/609/CEE) and approval of the local ethics committee (agreement D13-055-8). After 6 DIV, neurons were fixed, processed and imaged as previously described^2,68^. Neurons were fixed with 4% glutaraldehyde, 4% sucrose in PEM buffer (80mM PIPES pH 6.8, 5mM EGTA, 2mM MgCl_2_) for 10 minutes at room temperature, blocked in blocking buffer (phosphate buffer 0.1M with 0.22% gelatin, 0.1% Triton X-100) for 2h, stained with primary antibodies overnight, and then secondary antibodies for 1h in blocking buffer. Coverslips were placed in STORM buffer (Abbelight) and imaged on an N-STORM microscope (Nikon Instruments). A series of 60,000 images (67Hz frame rate) was acquired at full power with a 647nm laser, with progressive reactivation with a 405nm laser. Sequences of images were processed for localizations using the N-STORM software and 2D projections of the 3D-STORM data were generated using the ThunderSTORM plugin for ImageJ^69^ and the ChriSTORM scripts (http://github.com/cleterrier/ChriSTORM).

### SIM Super-resolution microscopy

A Nikon-SIM S microscope was used to acquire SIM images at two distinct locations along the axon of transfected neurons: proximal axon (around ∼100 μm from the cell body) and distal axon (∼200-300 μm from the cell body). Exposure times and laser powers were: 488nm, ∼200 msec at ∼80 percent laser power; 561 nm, ∼100 msec at ∼30 percent laser power. The objective was a CFI Apochromat TIRF 100XC Oil (NA 1.49) objective and the system was equipped with a TI2-FT N-SIM 405/488/561/640 quad band filter cube and corresponding emission filters. Images were acquired in 3D-SIM mode (15 raw images per frame) as Z-stacks with a spacing of 0.12 μm. The microscope used an ORCA-Fusion BT camera (Hamamatsu Photonics K.K. – C15440-20UP) and was controlled by NIS Elements 5.30.05 software. Reconstruction was performed using the N-SIM module within NIS-Elements. The acquired image pixel size was 0.065 μm and the final pixel size of processed SIM images was 0.032 μm, with an XYZ resolution of ∼120 x 120 x 240 nm.

### SIM image analysis

A set of macros (adapted from https://github.com/cleterrier/Process_Images) within Fiji^70^ was used to process the reconstructed SIM images. In brief, nd2 Z-stacks are initially extracted to tif files, then projected along Z using a maximum projection and all images from a given experiment are grouped as a hyperstack that was used for the analysis of axonal periodicity. A set of macros (adapted from https://github.com/cleterrier/Measure_ROIs) was used to classify periodic regions along axonal segments. Axons were manually traced along their whole length using the ‘segmented lines’ tool (line width of 10 pixels), and the visibly periodic segments along the axons were traced separately. The length of the “whole” and “periodic” line ROIs was then measured and used to calculate the % periodicity along axons for the different locations (proximal, distal) and conditions.

### Axonal FRAP

Neurons were transfected with the indicated fluorescent constructs at 5DIV with Lipofectamine 2000 (Invitrogen). At 7DIV, neurons were transferred to Hibernate media (Brainbits) supplemented with 2% B27, 2mM GlutaMAX, 0.4% D-glucose, 37.5 mM NaCl (HELF^66,71,72^) and maintained at 37°C for the duration of experiments (heated stage chamber, model STEV; World Precision Instrument, Inc.). Cells were imaged on the microscope described in Immunostaining section, which was also equipped with an Andor Mosaic3 Digital Micromirror Device (DMD) attached to a 405nm diode laser (450mW). Axons were identified by morphology, and only neurons with unambiguous morphology were selected for imaging^66,72^. Axonal segments of 1 μm length or full spectrin patches were bleached using the 405nm laser with 5 consecutive pulses of 1s each and imaged every 2 seconds for several minutes.

### Droplet FRAP

HEK293T cells were maintained in DMEM + GlutaMAX supplemented with 10% FBS and 1% penicillin-streptomycin. Cultures were transfected using Lipofectamine 2000 (Invitrogen) 18-20 hours prior to imaging in HELF media. Cells were imaged on a Leica DMi8 inverted confocal microscope equipped with a 100x oil-immersion objective, live cell imaging chamber, and white light laser for photobleaching. Droplets were imaged at 14.3fps (RGG-GFP-RGG), 8.33fps (βII-spectrin SR1-17; αII-spectrin full length protein + SR8-10; αII-spectrin full length protein + βII-spectrin SR1-17), 1.15fps (Adducin ΔN-term), or 4.4fps (all other constructs). For all FRAP experiments, droplets were imaged for 3 frames, bleached for one frame using 100% laser power at 405nm and 488nm, and then imaged for 600 frames. Bleaching regions were drawn individually for each droplet and either encompassed the entire droplet (full-droplet FRAP) or approximately 50% of the droplet area (partial droplet FRAP).

### FRAP quantification

Full droplet and axonal FRAP data were analyzed as previously described^73–75^. Background fluorescence values were subtracted from data at each time point, and photobleaching was corrected by normalizing data to an unbleached control region in the same image. Fluorescence curves were normalized between cells by subtracting the fluorescence immediately after bleaching (time = 0), and then dividing by the average of the 3-5 prebleach fluorescence time points. Fluorescence recovery rate and mobile fraction were calculated from an inverse exponential decay (*F(t) = A * (1 - e^-τt^)*, where *F* is fluorescence, *A* is the recovery plateau fluorescence, τ is the recovery time constant, and *t* is time) fit to the fluorescence recovery curve for each droplet. Fluorescence recovery curves were fit to the data with the OpenSolver add-in of Microsoft Excel 2013^76^ using the NOMAD nonlinear formula algorithm.

Partial droplet FRAP dip depth was calculated from the average of the lowest 10 data points in the unbleached half, excluding the first 5 frames after bleach. The internal diffusion rate (*D*) was determined as described previously^32^. We calculated the recovery curve within the droplet (*F*_diff_) as the difference between the recovery of the bleached half (*F*_half_), which relies on diffusion within the droplet and flux at the condensate boundary, and the recovery of the full droplet (*F*_full_), which only relies on boundary flux:

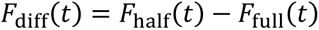

We then fit this curve to a simple diffusion model:

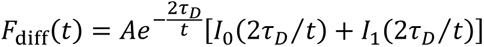

where *I*_0_ and *I*_1_ are modified Bessel functions of the 0^th^ and 1^st^ order, respectively, and τ*_D_* = *R*^2^/(4*D*), with *R* defined as the radius of the bleached region and *D* the internal diffusion coefficient.

### Protein Disorder Prediction

Protein disorder was predicted using the IUPred2A algorithm^77,78^ and cross-referenced with the D2P2 protein disorder prediction database, which aggregates disorder predictions from several additional algorithms^79^. A region was considered to be disordered if it scored above 0.5 according to IUPred2A and met consensus with at least seven of the nine predictors included in the D2P2 database. The protein sequences used for prediction were murine αII-spectrin (Sptan1, P16546-1), murine βII-spectrin (Sptbn1, Q62261-1), and murine α-adducin (Add1, Q9QYC0-1).

### Statistical analysis

Statistical tests were performed using Graphpad Prism. Normality of data sets was tested using the Shapiro-Wilk W test with α set at 0.05. Pairwise comparisons were made using a Student’s t test when both groups compared were found to be normally distributed: in other cases, a Mann-Whitney nonparametric U test was used. When more than two groups were compared, a one-way ANOVA or Kruskal-Wallis test was used in the case of normally or non-normally distributed data, respectively, with Dunn’s post-hoc pairwise comparisons against control groups when multiple comparison p was less than 0.05. Axonal FRAP curves were compared by two-way ANOVA interaction factor between time after bleach and condition. Actin colocalization with spectrin was compared between groups by two-way ANOVA interaction factor between axon segment and actin-containing fraction, with correction for multiple comparisons using the Bonferroni method. No *pre-hoc* sample size estimates were determined prior to experiments due to a lack of information on expected effect sizes. All data are presented as mean ± SEM. In all figures, significance is represented as follows: n.s., p > 0.05; *, p < 0.05; **, p < 0.01; ***, p < 0.001.

## SUPPLEMENTAL INFORMATION

**Supplementary Figure 1:**
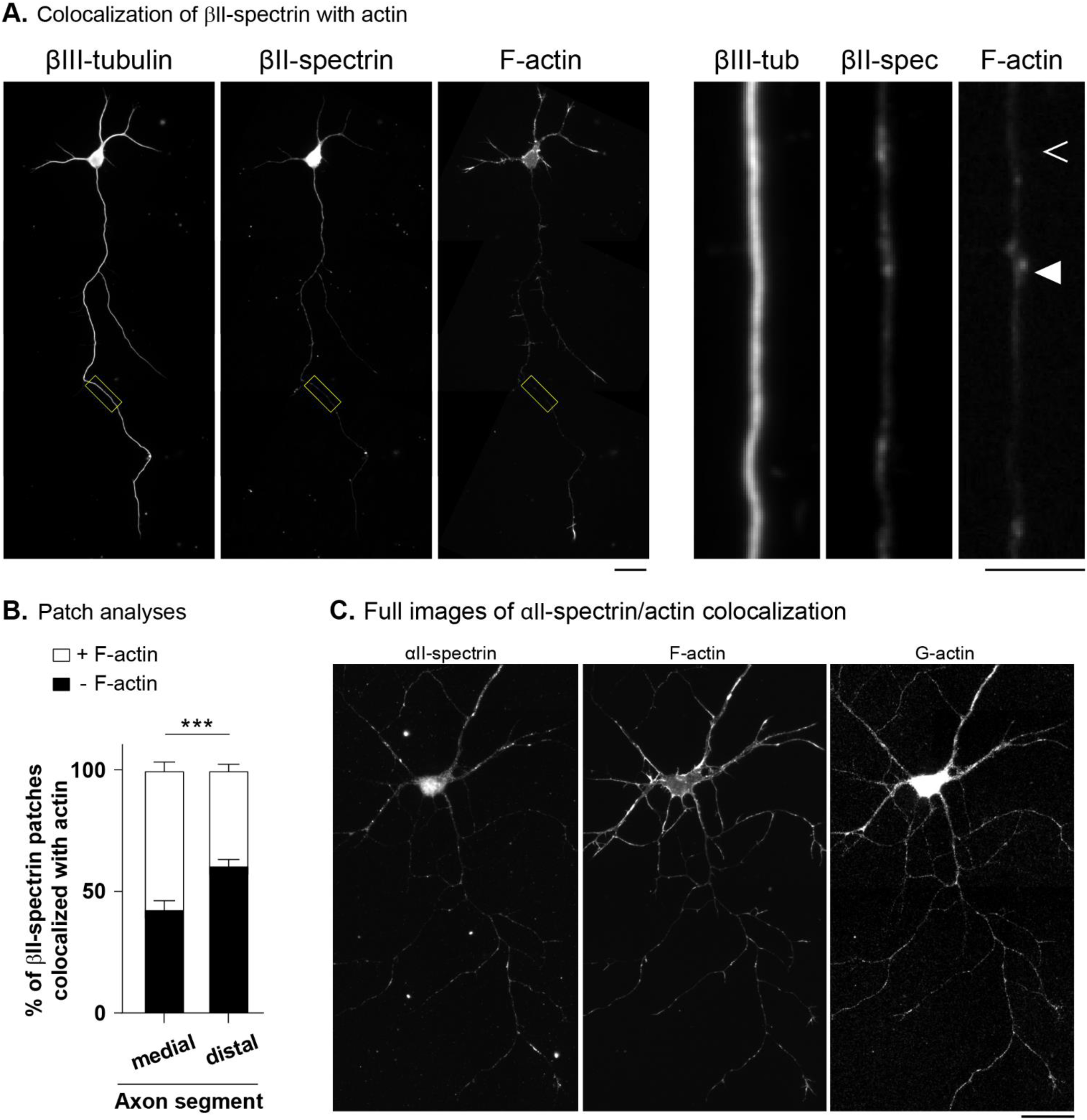
Distal spectrin patches have scant F-actin. **(A)** Cultured mouse hippocampal neurons were fixed at DIV-7 and stained with phalloidin (F-actin), as well as antibodies against βIII-tubulin and βII-spectrin. Note that the distal axon (yellow boxes with zoomed images on right) contains patches of spectrin, as well as foci of F-actin. Some patches of spectrin contain F-actin (filled arrowhead) while others do not (empty arrowhead). Scale bars = 10 µm (images on left), 2 µm (images on right). **(B)** Quantification of βII-spectrin patches colocalized with F-actin. Note that the number of spectrin patches colocalizing with F-actin is lower in the most distal parts of the axons (DIV-7 neurons; n=31 cells from 3 independent cultures; two-way ANOVA, actin fraction *x* axon position interaction ***, p < 0.001). **(C)** Staining in neurons related to Fig. 2A**-C**. Cultured mouse hippocampal neurons were fixed at DIV7, and stained with phalloidin (F-actin), DNaseI (G-actin) and an antibody against αII-spectrin. Scale bar = 10 µm.

**Supplementary Figure 2:**
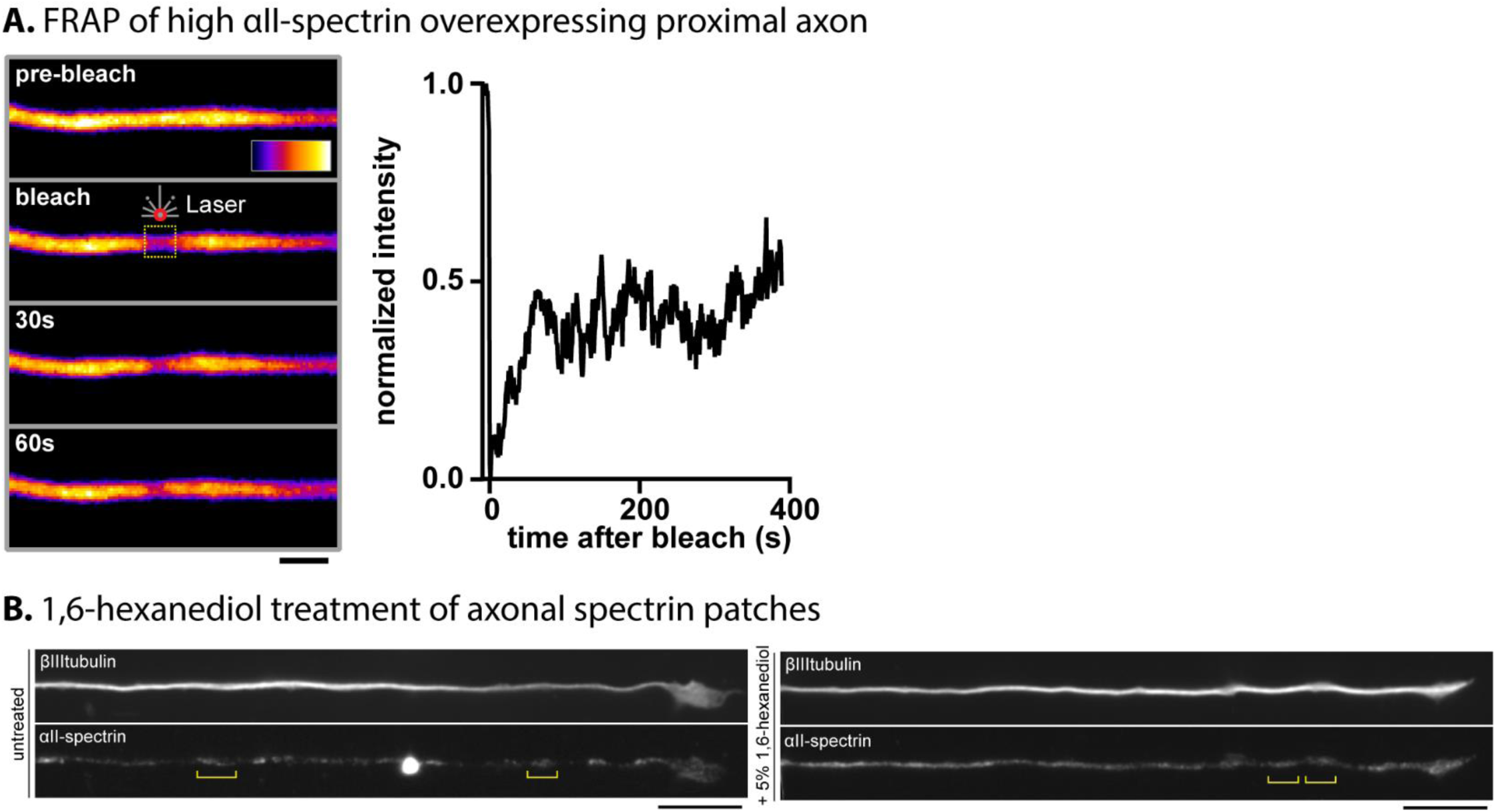
Full SIM images and hexanediol experiments. **(A)** Cultured hippocampal neurons were transfected with full-length αII-spectrin at DIV5 and imaged at DIV7 as in Fig.2. Axons were excluded from further imaging and analysis if proximal spectrin recovered quickly after photobleaching (see Zhong et al., 201421). Scale bar = 2 μm. **(B)** Primary mouse hippocampal neurons were treated with naïve media or media containing 5% 1,6-hexanediol for 30 minutes prior to fixation with 4% PFA and staining with antibodies against βIII-tubulin and αII-spectrin. Patches remain distinct in the distal axon (yellow brackets) despite hexanediol treatment, suggesting that patches are not reliant on weak, hydrophobic interactions. Scale bars = 10 µm.

**Supplementary Figure 3:**
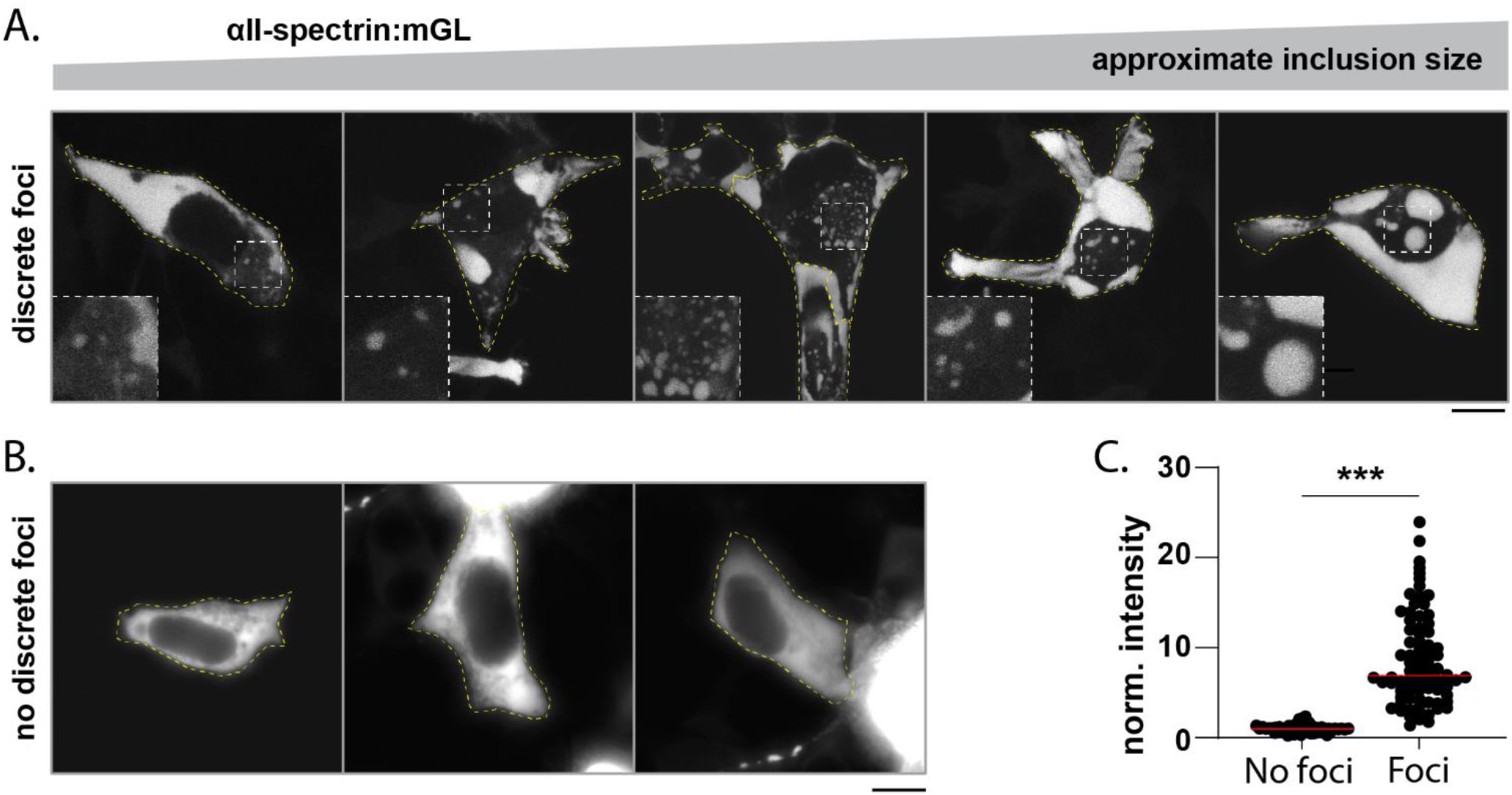
Overexpression of FL-αII-spectrin in in HEK293T cells leads to inclusions of variable shapes and sizes. **(A)** Fluorescent inclusions were seen in ∼ 59% of HEK293T cells overexpressing FL-αII-spectrin:mGL. Note that the inclusions had a range of sizes (insets), and there were also larger “sheet-like” accumulations in the cytoplasm. Scale bar = 10 µm. **(B)** Examples of cells expressing FL-αII-spectrin:mGLthat had no visible fluorescent foci. Scale bar = 10 µm. **(C)** Quantification of αII-spectrin:mGL fluorescence in cells with and without discrete foci. Fluorescence intensity in each frame was normalized to the average intensity of all cells without discrete foci. Cells with discrete foci had significantly more fluorescence intensity than those without foci, analogous to phase transitions with increasing concentrations of proteins *in vitro*. Red lines show median of each group. Data from 135 cells from 4 independent cultures. ***, p < 0.001.

**Supplementary Figure 4:**
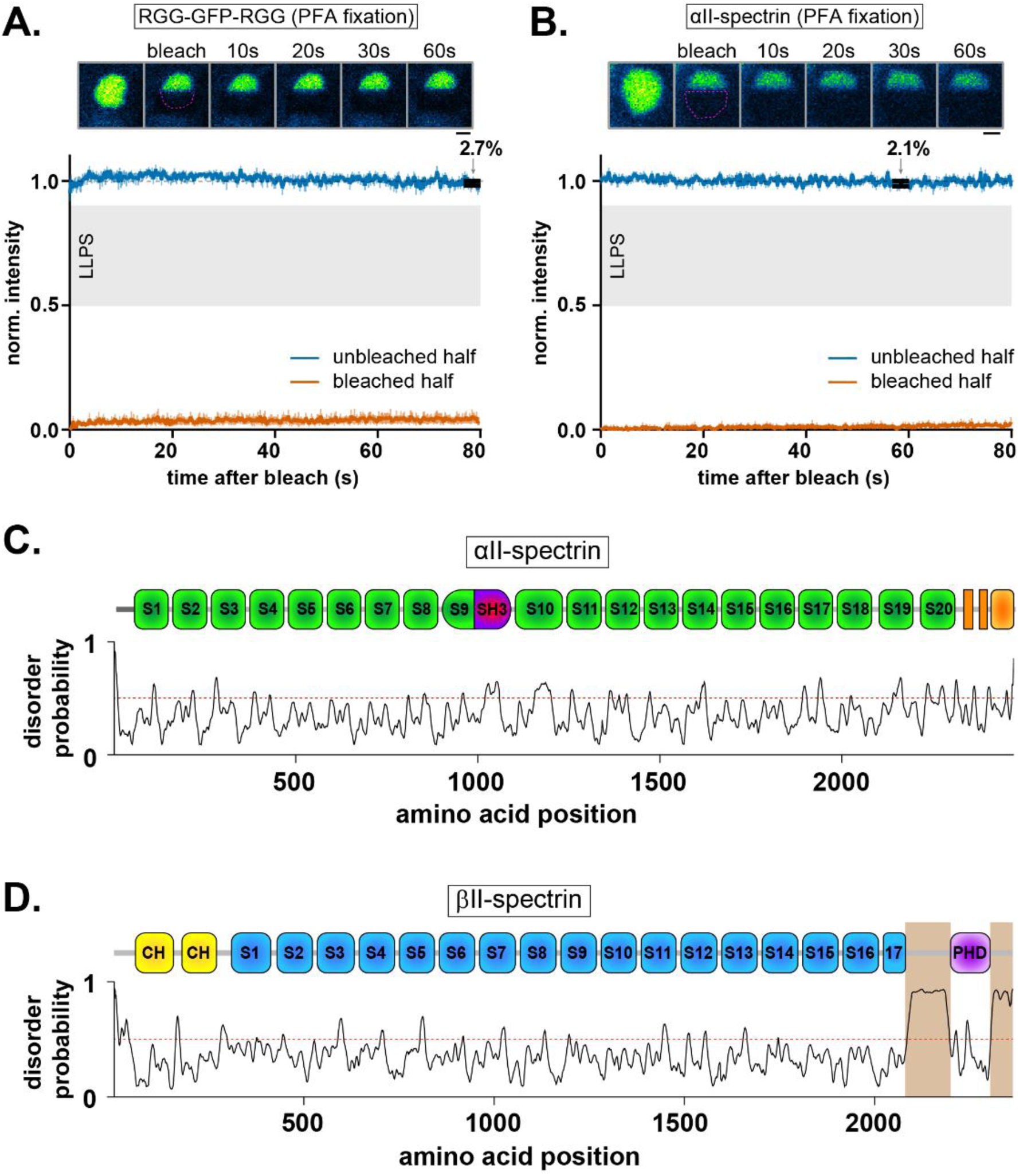
Additional controls for MOCHA-FRAP experiments and disorder predictions for αII/βII-spectrins. **(A)** MOCHA-FRAP assays of inclusions formed by RGG-GFP-RGG following 15 minutes of fixation with 4% PFA. Note that as expected, there is no redistribution of fluorescence after bleaching. Dip depth labeled at minimum fluorescence in the unbleached half (n=7 droplets, cells obtained from 3 independent cultures, scale bar = 1 µm). **(B)** MOCHA-FRAP assays of αII-spectrin:mGL inclusions following 15 minutes of fixation with 4% PFA, with no apparent redistribution of fluorescence after bleach. Dip depth labeled at minimum fluorescence in the unbleached half (n=7 droplets, cells obtained from 3 independent cultures, scale bar = 1 µm). **(C)** IUPred2A disorder-prediction of murine αII-spectrin sequence alongside a schematic of the protein with spectrin repeats 1-20 in green, SH3 domain in magenta, and calcium-binding EF hands in orange. While short stretches pass above the prediction threshold of 0.5, there are no clear intrinsically-disordered regions in the protein. **(D)** IUPred2A disorder-prediction of murine βII-spectrin sequence alongside a schematic of the protein showing spectrin repeats 1-17 in blue, actin-binding calponin homology (CH) domains in yellow, and membrane-binding pleckstrin homology domain (PHD) in purple. Regions of the protein in which disorder probability fell above the predictive threshold of 0.5 are indicated in tan highlight, spanning the full C-terminus following spectrin repeat 17 except for the PHD-domain.

**Supplementary Figure 5:**
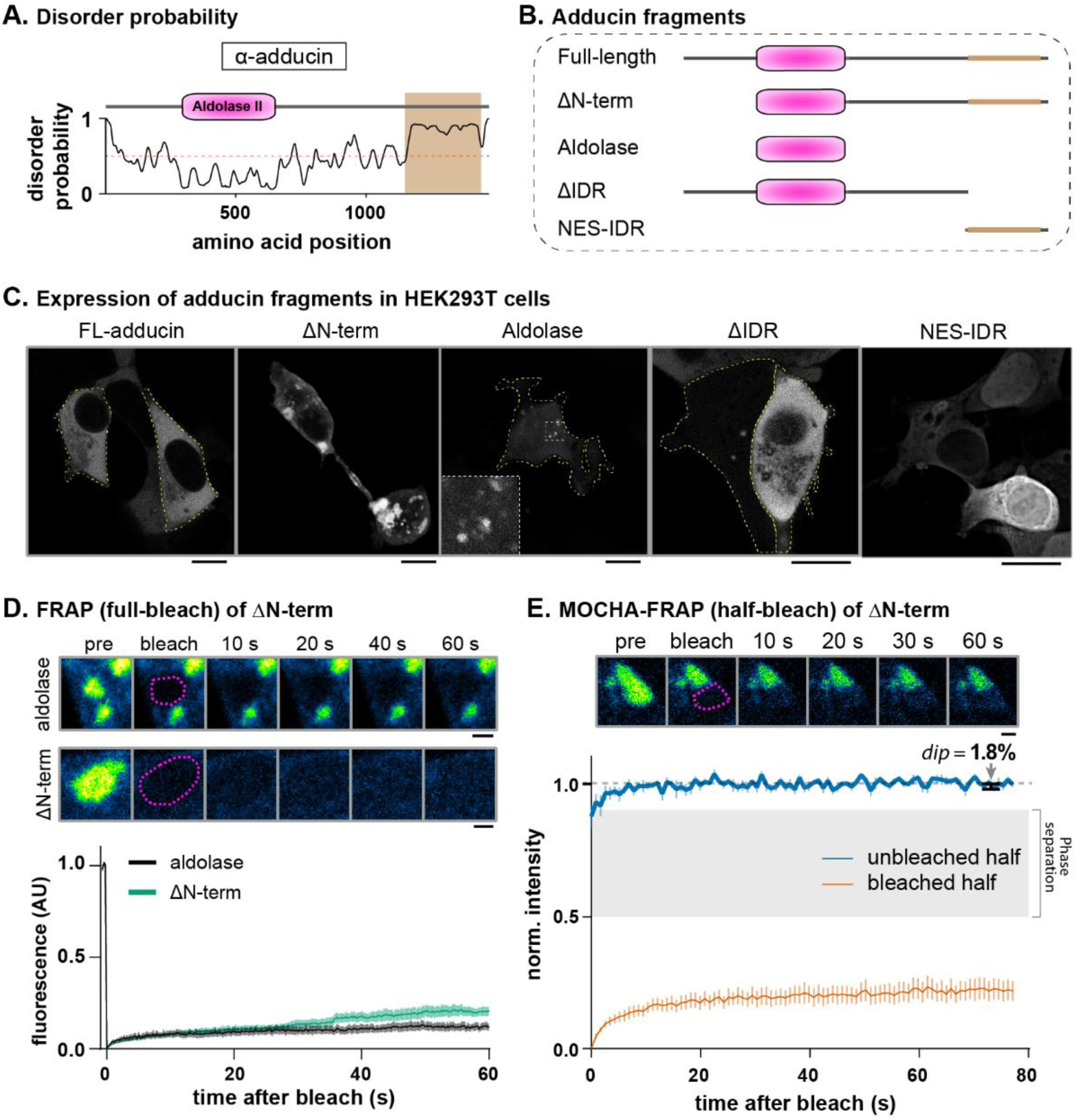
Expression of FL and domain-deletion adducin constructs. **(A)** IUPred2A disorder-prediction results from murine α-adducin sequence alongside a schematic of the protein showing aldolase II domain in pink. A region of the C-terminus of the protein falling above the predictive threshold of 0.5 is highlighted in tan. **(B)** Domain-deletion constructs of α-adducin that were overexpressed in HEK293T cells. Note that domains were chosen to either include or exclude the aldolase II domain or the intrinsically disordered region (IDR), which are the two main features of this protein. A predicted nuclear export signal (NES) is also present at the C-terminus. **(C)** Representative images showing overexpression of mGL-tagged FL and domain-deletion adducin constructs in HEK293T cells. While the distribution of FL-adducin:mGL is diffuse throughout the cytoplasm, ΔN-term induced large, bright inclusions in over half of the cells, while a construct containing only the aldolase-domain formed much smaller inclusions in all cells. Constructs lacking (or containing) only the IDR-domain was diffuse in all cells. Scale bars = 10 µm. **(D)** Full-bleach FRAP experiments of inclusions formed by ΔN-term and Aldolase adducin constructs showed minimal fluorescence recovery after bleaching (n = 5-7, in cells from 3 independent experiments; scale bars = 1 µm). **(E)** MOCHA-FRAP analyses of inclusions formed by ΔN-term construct showing minimal overall recovery of fluorescence as well as no exchange between unbleached and bleached halves. Dip depth shown in black at minimum fluorescence following the first 5 frames (n= 7, in cells from 3 independent experiments; scale bars = 1 µm).

**Supplementary Figure 6:**
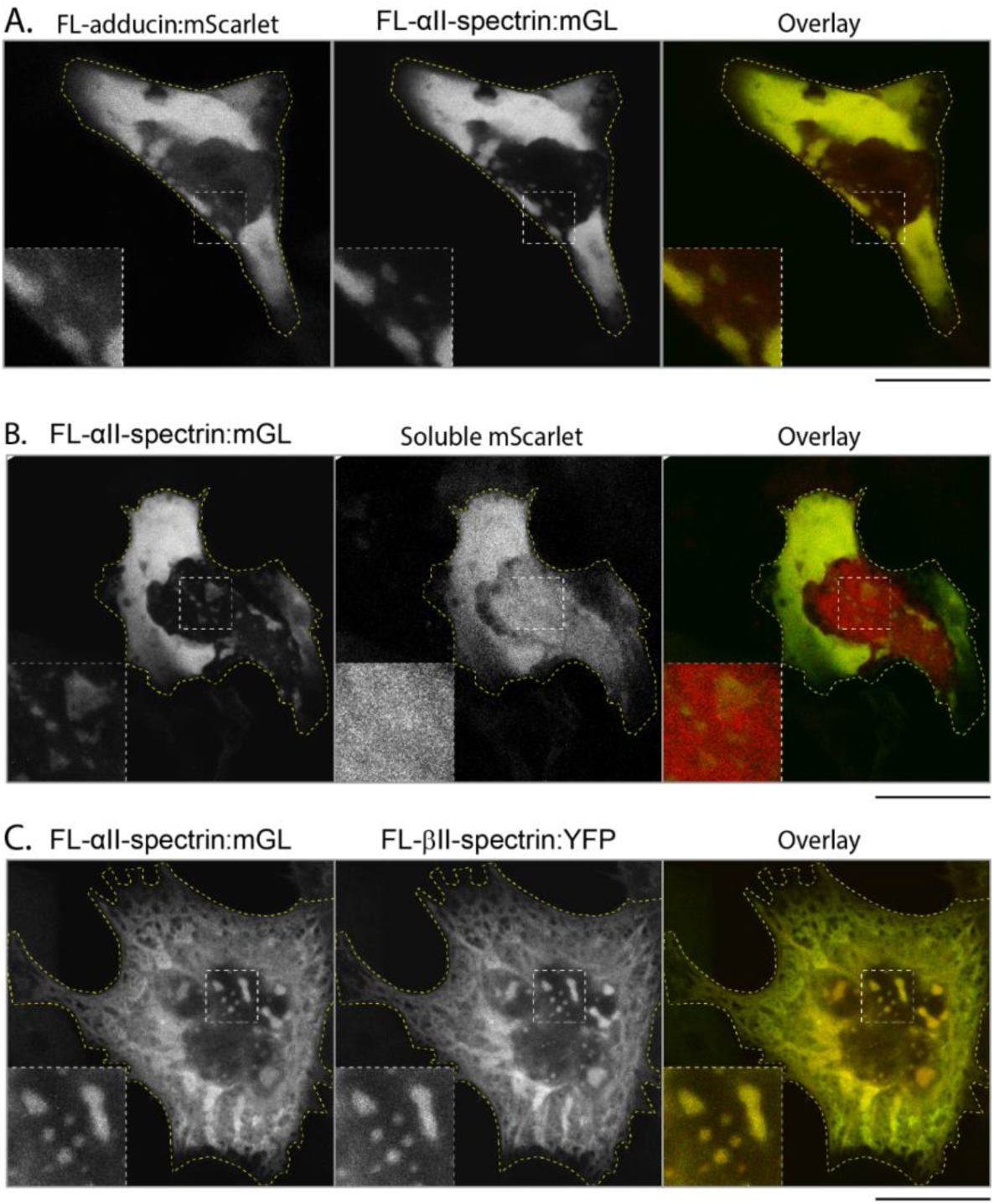
Adducin and βII-spectrin are incorporated into αII-spectrin droplets. **(A)** HEK293T cells co-transfected with plasmids encoding FL-α-adducin:mScarlet and αII-spectrin:mGL which colocalize in rounded inclusions reminiscent of those formed by αII-spectrin alone (white dashed outline inset). Cell border delineated with yellow dashed outline. Scale bar = 10 µm. **(B)** HEK293T cells co-transfected with plasmids encoding FL-αII-spectrin:mGL and soluble mScarlet. Spectrin forms rounded inclusions as when expressed alone, while mScarlet remains primarily diffuse and is not enriched at the location of spectrin droplets (white dashed outline inset). Cell border delineated with yellow dashed outline. Scale bar = 10 µm. **(C)** HEK293T cells co-transfected with plasmids encoding FL-αII-spectrin:mGL and βII-spectrin:YFP which colocalize in rounded inclusions (white dashed outline inset) as well as broad areas with a filamentous appearance in the rest of the cell. Cell border delineated with yellow dashed outline. Scale bar = 10 µm.

### Supplementary Movie legends

**<u>Supplemental movie 1</u>: αII-spectrin:mGL patches in distal axons.** Timelapse images of spectrin patches within 50 μm of the axonal growth cone (time interval between frames = 2 s, >18 minutes total imaging time). Note that patches are largely immobile over the course of imaging. Scale bar = 5 μm.

**<u>Supplemental movie 2</u>: Z-stack of HEK293T cells expressing FL-βII-spectrin:mGL.** Note membranous enrichment in addition to cytosolic pool. Scale bar = 5 μm.

**<u>Supplemental movie 3</u>: Timelapse imaging of αII-spectrin:mGL inclusions in HEK293T cells.** Timelapse images αII-spectrin:mGL inclusions in HEK293T cells approximately 20 hours after transfection. (time interval between frames = 1 s, 2.5 minutes total imaging time). Note that the inclusions are mostly immobile, and do not show splitting/merging behaviors. Scale bar = 2 μm.

